# Retrograde signalling mediates cellular adaptation to mitochondrial DNA copy number alterations

**DOI:** 10.64898/2026.04.22.720057

**Authors:** Alissa Benedikt, Kim Job, Xiaoping Li, Soumodeep Sarkar, Luisa Hernández Götz, Igor V. Kukhtevich, Francesco Padovani, Yagya Chadha, Jurgita Paukštytė, Juha Saarikangas, Jan M. Skotheim, Robert Schneider, Matthew P. Swaffer, Antonio Scialdone, Michael C. Lanz, Andreas Kohler, Kurt M. Schmoller

**Affiliations:** Institute of Functional Epigenetics, Molecular Targets and Therapeutics Center, Helmholtz Zentrum München, Neuherberg, Germany; Institute of Epigenetics and Stem Cells, Stem Cell Center, Helmholtz Zentrum München, Munich, Germany; Institute of Computational Biology, Computational Health Center, Helmholtz Zentrum München, Neuherberg, Germany; Department of Medical Biochemistry and Biophysics, Umeå University, Umeå, Sweden; Centre for Cell Biology, University of Edinburgh, Edinburgh, UK; Helsinki Institute of Life Science HiLIFE, and the Faculty of Biological and Environmental Sciences, University of Helsinki, Helsinki, Finland; Department of Biology, Stanford University, California, US

## Abstract

Eukaryotic cells maintain multiple copies of the mitochondrial genome, which is essential for cellular metabolism. Accordingly, alterations in the mitochondrial DNA (mtDNA) copy number are associated with severe human diseases and ageing. However, the mechanisms through which cells regulate mtDNA copy number and the cellular consequences of altered copy number remain poorly understood. Here, using budding yeast as a model, we show that mtDNA copy number is determined by the amount of three limiting factors, Mip1, Abf2, and Rim1. By synthetically tuning the concentrations of only these three proteins, we can modulate mtDNA dosage inside the cell. This revealed that cells are surprisingly robust to mtDNA copy number alterations, with increased copy numbers even accelerating cell growth. Our findings suggest that this robustness is due to protein dosage compensation and independent regulation of mitochondrial morphology. Mechanistically, we identified a critical role of the retrograde signalling pathway for this adaptation. We show that signalling from mitochondria to the nucleus is upregulated in cells with higher mtDNA copy number, and disruption of this regulation diminishes their faster growth. Taken together, our work reveals regulatory principles that enable cells to adapt to mtDNA copy number alterations.

## Introduction

Mitochondria are essential organelles in almost all eukaryotic cells. Reminiscent of their endosymbiotic origin, they contain their own mitochondrial genome. This mitochondrial DNA (mtDNA) typically encodes for several mitochondrial rRNAs, tRNAs, and proteins, in particular subunits of the oxidative phosphorylation (OXPHOS) complexes^1–4^. In most organisms, including yeast and mammals, cells contain multiple copies of mtDNA. The copy number can vary drastically not only between species, but also between cell types, during development, and under different environmental conditions^5–8^. Moreover, mtDNA copy number changes are associated with human ageing^9–11^, as well as neurodegenerative diseases such as Alzheimer’s and Parkinson’s disease^12–15^. Both up-and downregulation of mtDNA copy number have also been observed in various types of cancer, where they have been linked to mutational load, disease progression, and therapy response^16–19^. However, these relationships, and even their causality, can be highly specific to cancer type and tissue^17,20,21^, which makes a mechanistic interpretation difficult. Even on a cellular level, it is unclear how mtDNA copy number changes translate into functional consequences. Thus, despite the obvious importance of mtDNA copy number for human health, the mechanistic consequences of copy number alterations are surprisingly poorly understood.

One reason for our poor understanding of the cellular response to mtDNA copy number changes is that, so far, we have lacked suitable models to investigate them. Developing such a model requires understanding the mechanism by which mtDNA copy number is regulated in the cell. Then, targeted manipulation of this mechanism can generate controlled copy number variation whose effects can be studied. In budding yeast, it has recently been shown that, within a given nutrient condition, mtDNA copy number increases in proportion to cell volume, ensuring constant mtDNA concentrations throughout the cell cycle^5^. This coupling of mtDNA copy number to cell volume is controlled by the dosage of limiting nuclear-encoded machinery for mtDNA replication and maintenance, which, like most proteins, increases in abundance with cell volume^5^. Pointing towards a potential genetic tool to tune mtDNA copy number, two proteins, Mip1 and Abf2, have previously been identified as the major limiting factors^5^. Mip1 is the mtDNA polymerase^22^, and the TFAM-homolog Abf2 packages mtDNA in so-called nucleoids^23^, clusters of mtDNA that are regularly distributed in the mitochondrial network^24,25^. Specifically, reducing expression of *MIP1* and *ABF2* to 50% through hemizygous (HZ) deletions in diploid cells leads to a near twofold reduction of mtDNA copy number^5^. However, this picture is incomplete as a twofold overexpression of these factors in haploid cells does not result in the predicted twofold increase (Fig. 1a, data from Seel et al.^5^), suggesting that other key factors must also be dosage-limiting for mtDNA copy number.

**Figure 1:**
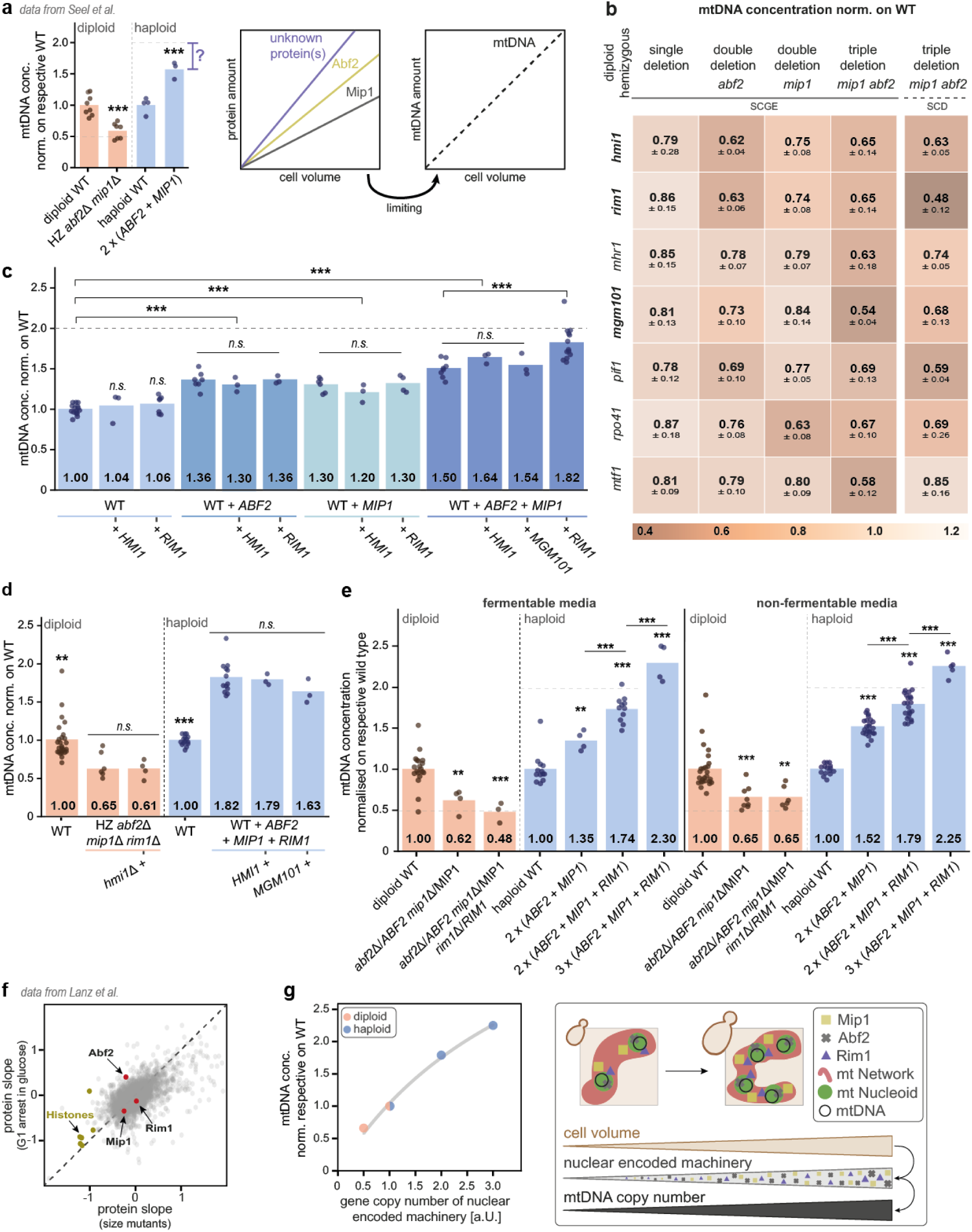
mtDNA copy number is coupled to cell volume by a nuclear-encoded limiting machinery composed of Abf2, Mip1, and Rim1. **a)** Protein amounts of Abf2 and Mip1 increase with cell volume, leading to increasing mtDNA copy numbers. Consistent with Abf2 and Mip1 being the major limiting factors for mtDNA maintenance, mtDNA copy number decreases to about 50% in a diploid strain with hemizygous deletions of both genes. However, the less than twofold increase in a haploid strain with additional copies of both genes hints towards additional proteins limiting mtDNA copy number. Data from Seel et al.^5^. **b)** mtDNA concentration normalised on diploid wild type (WT) for diploid hemizygous deletion strains determined by DNA-qPCR. Strains grown in non-fermentable media (SCGE; straight line) and triple deletions additionally grown in fermentable media (SCD; dashed line). Shown are means of at least three replicates per strain and standard deviations. **c)** mtDNA concentration normalised on haploid WT for strains with additional copies of one or multiple genes grown in non-fermentable media determined by DNA-qPCR. Shown are single replicates (blue dots) and the mean of all replicates (bars). Significance was determined by a two-tailed t-test (* p<0.05, ** p<0.01, *** p<0.001). **d)** mtDNA concentration normalised on respective WT for strains grown in non-fermentable media, determined by DNA-qPCR. Shown are single replicates (dots) and the mean of all replicates (bar) for diploids (orange) and haploids (blue). Significance was determined by a two-tailed t-test (* p<0.05, ** p<0.01, *** p<0.001). **e)** mtDNA concentration normalised on respective WT for various strains grown in fermentable (top) or non-fermentable media (bottom), determined by DNA-qPCR. Diploids (orange) include WT, hemizygous deletion of *ABF2* and *MIP1*, and hemizygous deletion of *ABF2, MIP1,* and *RIM1*; haploids (blue) include WT, strains with two copies of *ABF2* and *MIP1*, as well as with two or three copies of *ABF2, MIP1,* and *RIM1*. Shown are single replicates (dots) and the mean of all replicates (bars). Significance was determined by a two-tailed t-test (* p<0.05, ** p<0.01, *** p<0.001). **f)** Protein slopes characterising the cell volume-dependence for cells grown in glucose media were determined from G1 arrests as a function of those determined from size mutants. Histone (yellow) amounts are independent of cell volume, while Abf2, Mip1, and Rim1 (red) amounts increase with increasing cell volume. Data from Lanz et al^28^. **g)** mtDNA concentration normalised on respective WT increases with increasing gene copy number of the nuclear-encoded machinery (*ABF2*, *MIP1*, *RIM1*) (left). The grey line shows a Michaelis-Menten-like fit with a maximum of 5.4 and a Michaelis constant of 4.2. Illustration of mtDNA copy number regulation in yeast (right): increasing cell volumes lead to increasing amounts of the nuclear-encoded machinery, causing increased mtDNA copy numbers.

Here, we identify the single-strand DNA-binding protein Rim1 as the third limiting factor for mtDNA maintenance. Across different nutrient conditions, mtDNA copy number follows the abundance of the limiting machinery, consisting of Abf2, Mip1, and Rim1, over a severalfold range. By modulating the dosage of this limiting machinery, we are able to alter mtDNA concentrations across a fourfold range – providing a powerful system to investigate the consequences on cell function. Using this system, we find that cells are surprisingly robust towards alterations in mtDNA copy number, and that increased mtDNA concentrations even lead to faster cell growth. We attribute this robustness to an mtDNA-independent regulation of mitochondrial morphology, as well as dosage compensation of mitochondrially encoded proteins. We identified the retrograde signalling pathway, which triggers a nuclear response to mitochondrial changes, as an essential factor underlying the observed adaptation. Cells with higher mtDNA copy numbers show increased retrograde signalling activity, and disrupting the retrograde pathway diminishes their faster growth.

## Results

### Mip1, Abf2, and Rim1 are the major limiting factors for mtDNA copy number

To identify the factors that limit mtDNA copy number in addition to Mip1 and Abf2, we first created a series of hemizygous deletions in four different diploid strains: a wild type, a hemizygous deletion of *ABF2* (*abf2*Δ*/ABF2*), a hemizygous deletion of *MIP1* (*mip1*Δ*/MIP1*), and a hemizygous double deletion (*abf2*Δ*/ABF2, mip1*Δ*/MIP1*). In each of these strains, we deleted one allele of one of seven genes of interest, which we selected because they individually show small effects on mtDNA copy numbers^5^. Measuring mtDNA to nuclear DNA (nDNA) ratio, bud fraction, and cell volume, we determined the mtDNA concentration when cells were grown on either fermentable (SCD, synthetic complete media with 2% glucose) or non-fermentable (SCGE, synthetic complete media with 2% glycerol and 1% ethanol) carbon sources.

We found that the hemizygous triple deletions of *MIP1*, *ABF2,* and *RIM1,* or of *MIP1*, *ABF2,* and *MGM101,* lead to the lowest mtDNA concentrations for growth on SCD and SCGE media, respectively (Fig. 1b; Fig S1a). Moreover, across the four background strains and two different growth media, deleting *HMI1* showed the most consistent reduction among the candidate genes. We therefore decided to proceed with *RIM1*, *MGM101,* and *HMI1*, and test dosage limitation further by adding an additional copy of each of these genes into each of the following haploid strains: wild type, a strain containing one extra copy of *ABF2* (2x *ABF2*), a strain containing one extra copy of *MIP1* (2x *MIP1*), or a strain containing extra copies of both *ABF2* and *MIP1* (2x (*ABF2*+*MIP1*)) Only the addition of an extra copy of *RIM1* led to a significant increase in mtDNA copy number in the strain that already carries additional copies of *ABF2* and *MIP1* (Fig. 1c; Fig S1b-c). Since we found no further effect when adding extra copies or deleting one allele of either *HMI1* or *MGM101* in the *ABF2*, *MIP1,* and *RIM1* triple addition or hemizygous deletions strains, respectively (Fig. 1d; Fig. S1d), we concluded that the ssDNA-binding protein Rim1^26^ is the only major limiting factor for mtDNA copy number along with the mtDNA packaging factor Abf2^27^ and mtDNA polymerase Mip1^22^.

Consistent with this, the simultaneous threefold overexpression of *ABF2*, *MIP1,* and *RIM1,* leads to a further increase of the mtDNA concentration to 225% compared to the wild type (Fig. 1e). This also exceeds the concentration that results from stronger overexpression of any of the individual factors, which we achieved using hormone-inducible promoters (for *ABF2* and *MIP1*) or multiple copy insertion (for *RIM1*) (Fig. S1e-g). This suggests that, in wild-type cells, the concentrations of the three factors are balanced such that increasing any of them alone simply renders the others limiting. Only when all three factors are manipulated together, mtDNA concentration increases almost proportionally with the concentration of the limiting machinery, allowing us to manipulate mtDNA concentrations over a roughly fourfold range centred around the wild type. Although mtDNA concentration varies substantially depending on the carbon source, the relative change caused by manipulating the limiting machinery remains similar (Fig. S1h). As for *MIP1* and *ABF2*, expression of *RIM1* also increases with cell volume (Fig. 1f). Our results therefore suggest that Mip1, Abf2, and Rim1 constitute the limiting machinery for mtDNA maintenance, whose abundance increases in proportion to cell volume and thereby directly couples mtDNA copy number to cell volume (Fig. 1g).

### Increased mtDNA copy number leads to reduced petite frequency and faster growth

Having established a system to manipulate mtDNA concentration via the key limiting factors (Fig. 1g), we next investigated the consequences of mtDNA copy number changes on cell function.

We found that the mtDNA copy number changes are not accompanied by any substantial changes in cell volume (Fig. 2a; Fig S2a) or budding behaviour (Fig. 2b; Fig S2b). Next, we estimated steady-state doubling times, which suggested that in non-fermentable media, the hemizygous deletions grow more slowly than the wild type, whereas additional copies of the limiting machinery lead to faster growth (Fig. 2c). To confirm this result, we performed competition assays for the haploid strains with increased mtDNA copy number. To exclude possible contributions from the auxotrophic markers, we compared each strain with a suitable control strain. Consistent with the growth rate measurements, we found that starting from initially 25% in the mixed culture, all three tested strains with higher mtDNA concentrations outcompeted their corresponding control strains within 100-150 hours (Fig. 2d; Fig. S2c-d). Taken together, our findings suggest that increasing mtDNA concentrations results in slightly faster growth on non-fermentable media.

**Figure 2:**
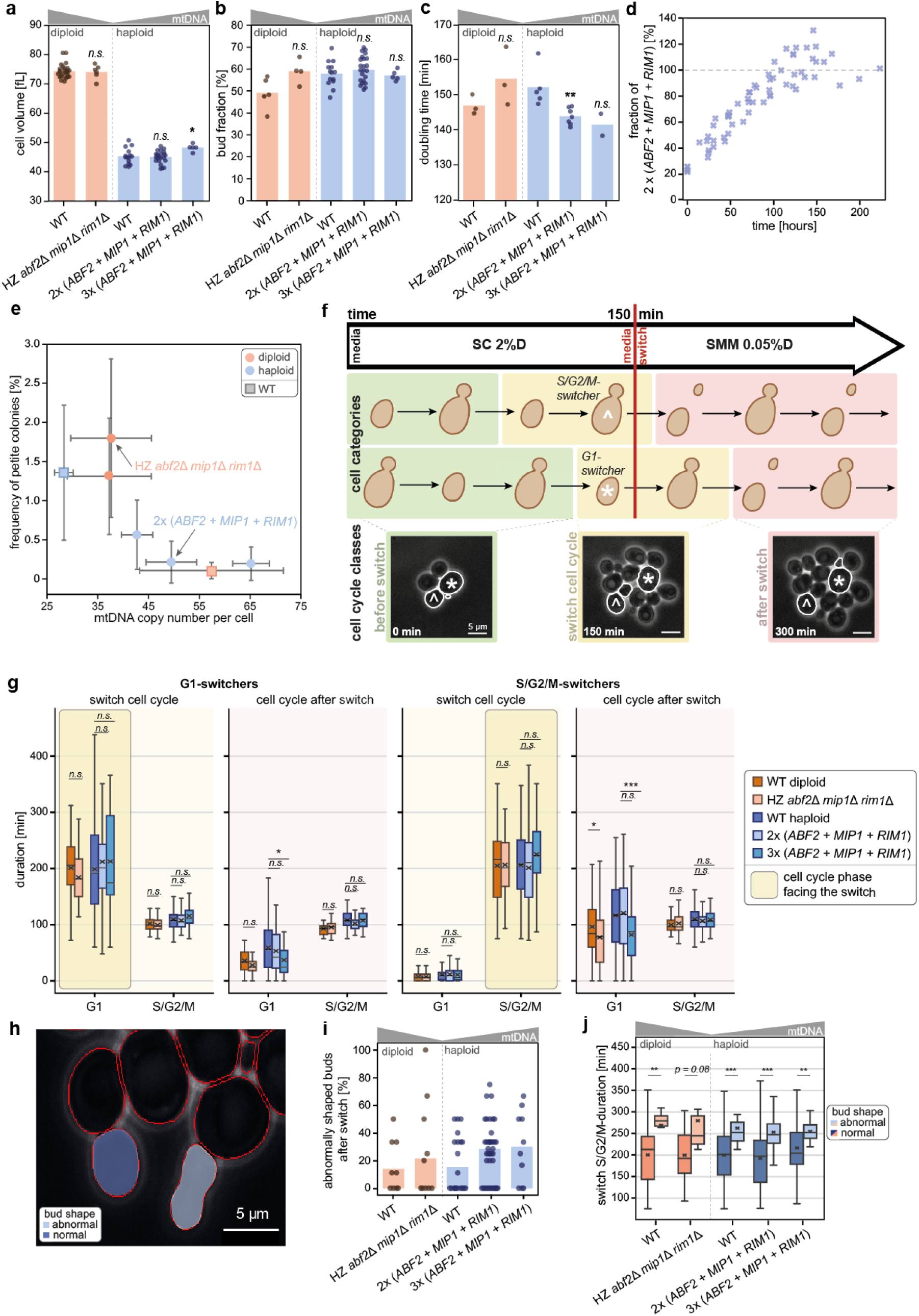
Cells are robust to mtDNA copy number changes across a fourfold range, and increased copy number leads to faster growth in non-fermentable media. **a-c)** Cell physiology measurements for various strains grown in non-fermentable media. Diploids (orange) include wild type (WT), and a hemizygous deletion of *ABF2, MIP1,* and *RIM1*; haploids (blue) include WT, and strains with two or three copies of *ABF2, MIP1,* and *RIM1*. Three independent clones were measured and pooled for the strains with two copies of the three factors. Shown are single replicates (dots) and the mean of all replicates (bars). Significance was determined by a two-tailed t-test (* p<0.05, ** p<0.01, *** p<0.001). **a)** Mean cell volume determined by Coulter counter; **b)** Percentage of budding cells determined by visual inspection under the microscope; **c)** Steady-state doubling time determined by OD_600_-measurements. **d)** Competition assay testing the growth of a strain with two copies of *ABF2, MIP1,* and *RIM1* against an auxotrophy-corrected control strain. The fraction of the two-copy strain in the mixture was initially 25%, and its dynamic development was monitored by DNA-qPCR. Crosses show individual measurements from three individual and pooled clones, each measured in three replicates. **e)** Percentage of petite colonies as a function of mtDNA copy number per cell for diploid (orange) and haploid (blue) strains. Both wild types shown as squares. Error bars indicate standard deviations. **f)** Illustration of media-switch time-lapse microscopy experiment using a custom microfluidic device: Initially, cells are grown in synthetic complete media (SC) with 2% glucose. 150 minutes after the start of the experiment, the media is switched to a minimal media (SMM) containing 0.05% glucose. Depending on the cell cycle phase in which they face the media switch, cells are categorised as either S/G2/M-switchers (^) or G1-switchers (*) (middle). Additionally, cell cycle classes are subdivided (bottom) into before the switch (green), switch cell cycle (yellow) and after the switch (red). Scale bars represent 5 µm. **g)** Phase durations are plotted for G1-switchers (2 left panels) and S/G2/M-switchers (2 right panels), highlighting the switch cell cycle phase and the cell cycle after the switch. Boxplots show the interquartile range (box), total range of data (whisker), median (line) and mean (cross). Significance was determined compared to the respective wild type by a two-tailed t-test (* p<0.05, ** p<0.01, *** p<0.001). Number of cells analysed for G1-switchers (left): n_WT diploid_ = 32 (dark orange), n_HZ_ *_abf2_*_Δ*mip1*Δ*rim1*Δ_ = 32 (light orange), n_WThaploid_ = 83 (dark blue), n_2x (*ABF2+MIP1+RIM1*)_ = 122 (light blue; three individual clones pooled), n_3x (*ABF2+MIP1+RIM1*)_ = 28 (turquoise), and for S/G2/M-switchers (right): n_WT diploid_ = 87 (dark orange), n_HZ_ *_abf2_*_Δ*mip1*Δ*rim1*Δ_ = 69 (light orange), n_WT haploid_ = 155 (dark blue), n_2x (*ABF2+MIP1+RIM1*)_ = 232 (light blue; three individual clones pooled), n_3x (*ABF2+MIP1+RIM1*)_ = 47 (turquoise). **h)** Example of normally-shaped bud (dark blue) and abnormal bud formation (light blue) for cells with two copies of *ABF2* and *MIP1*. Scale bar represents 5 µm. **i)** Percentage of abnormally shaped buds for S/G2/M-switchers whose buds are smaller than 26 fL at the time of the nutrient switch. Shown are single replicates (dots) and the mean of all replicates (bars). Number of buds analysed: n_WT diploid_ = 43, n_HZ *abf2*Δ_*_mip1_*_Δ_*_rim1_*_Δ_ = 28, n_WT haploid_ = 98, n_2x (*ABF2+MIP1+RIM1*)_ = 159 (three individual clones pooled), n_3x (*ABF2+MIP1+RIM1*)_ = 30. **j)** Duration of S/G2/M-phase in the switch cell cycle for normally (dark) and abnormally shaped buds (light), for diploid (orange) and haploid (blue) strains. Boxplots show the interquartile range (box), total range of data (whisker), median (line) and mean (cross). Significance was determined comparing the two types of bud shapes by a two-tailed t-test (* p<0.05, ** p<0.01, *** p<0.001).

Loss of functional mitochondrial DNA leads to the formation of petite colonies^29^, which are unable to perform respiration^30^. Previous studies have shown that increased mtDNA copy numbers in *cim1*Δ cells correlate with a decreased frequency of petite colonies^31^ and proposed that petite frequency negatively correlates with mtDNA copy number^32^. Additionally, each of the limiting factors Abf2^33^, Mip1^22,34–36^ and Rim1^26^ has previously been reported to influence the petite frequency. We therefore decided to test our strains to study the appearance of petite colonies as a function of mtDNA copy number. We showed that an increased mtDNA copy number leads to a decreased loss of respiratory capacity (Fig. 2e). In line with previous studies^37^, the haploid wild-type strain shows a higher frequency of petite colonies than the diploid wild-type strain. Interestingly, strains show comparable frequencies of petite colonies even across different ploidies if they have comparable mtDNA copy numbers, as for example the diploid hemizygous *abf2*, *mip1*, and *rim1* deletion strain compared to the haploid wild type, or the haploid triple addition (*MIP1*, *ABF2*, and *RIM*1) strain compared to the diploid wild type (Fig. 2e). This suggests that mtDNA copy number, rather than ploidy, is the major determinant of petite colony formation, consistent with stochastic loss of mtDNA at cell division being a major factor contributing to loss of mtDNA.

As increased mtDNA concentrations lead to faster cell growth and a reduced frequency with which cells lose respiratory capacity, we wondered whether increased mtDNA concentration would be detrimental under stress conditions. For this, we used spot assays to investigate the effect of mtDNA copy number changes on a variety of conditions, including replicative stress (100 nM hydroxyurea), osmotic stress (0.6 M sodium chloride), and different temperatures. We found no pronounced differences between strains with varying mtDNA copy number in any of the tested conditions (Figs. S3-S5).

### Cell cycle progression is surprisingly robust towards mtDNA copy number alterations

While bulk experiments provide robust measurements of average cell behaviour, they mask potential heterogeneity on the single-cell level. To address whether mtDNA copy number changes lead to more pronounced phenotypes in a subpopulation of cells, particularly with regard to cell cycle progression^38,39^, we used time-lapse live-cell microscopy. First, we investigated cells growing in a microfluidic device^40^ under steady-state conditions with a constant supply of non-fermentable media (SCGE). We analysed cell growth and cell cycle progression over multiple cell divisions using the software Cell-ACDC^41,42^. While this experiment confirmed that cells with increased mtDNA concentration grow faster, we did not observe any additional differences in cell cycle kinetics (Fig. S6a-b).

Finally, we wondered whether increased mtDNA copy numbers would be a disadvantage for cells when facing low glucose conditions. To test this, we imaged cells growing in steady-state on our standard fermentable media SCD (2% glucose) for 2 hours, and then switched to the synthetic minimal media SMMD with 0.05 % glucose (Fig. 2f). Consistent with bulk growth rate measurements in SCD media (Fig. S6c), mtDNA copy number does not significantly affect the doubling time during the initial steady-state growth (Fig S6d). After the media switch, cells show a temporary arrest in their current cell cycle phase^43^. Again, we did not observe any major differences between strains with varying mtDNA copy number (Fig. 2g; Fig. S7a). However, we noticed that the strains with altered mtDNA copy number showed an increased frequency of buds with abnormal morphology (Fig. 2h-i). This abnormal morphology was only observed in buds smaller than 26 fL at the time of the media switch. Cells that face the media switch in S/G2/M (“S/G2/M-switchers”) and develop this abnormal bud phenotype arrested significantly longer before proceeding to cell division (Fig. 2j). However, they divided at similar total cell volumes compared to cells with normally shaped buds (Fig. S7b). Thus, even under dynamic nutrient transitions, conditions that often reveal hidden sensitivities, cells remain largely robust to the mtDNA copy number variation.

In summary, we found that with increasing mtDNA concentration, budding yeast shows a lower frequency of losing respiratory capacity and faster growth in non-fermentable media. Notably, even under the diverse stress conditions examined, the phenotypic effects remain relatively subtle, indicating that cells are robust to mtDNA copy number variation within an approximately fourfold range around the wild-type level.

### Mitochondrial network morphology and nucleoid numbers show only a weak dependence on mtDNA copy number

Next, we asked how changes in mtDNA copy number affect the spatial organisation of the mitochondrial network and mtDNA-containing nucleoids. To visualise the mtDNA in cells growing either in fermentable or non-fermentable media, we used mt-Kaede-HI-NESS, a fluorescent reporter that is targeted to mitochondria and binds non-specifically to DNA but has a preference for AT-rich regions^44–46^ (Fig. 3a-b). In addition, we used mScarlet-I3 targeted to the mitochondrial matrix to visualise the mitochondrial network^47^. As expected^5^, we found that in each strain, network volume and nucleoid number both increase with cell volume (Fig. S8a). However, mtDNA copy number changes did not lead to substantial differences in the mitochondrial network volume, in either fermentable or non-fermentable media (Fig. 3c; Fig. S8b). We also did not observe a consistent mtDNA-dependency of mitochondrial network fragment number – only in fermentable media, we found that haploids with increased mtDNA copy numbers have a more connected mitochondrial network (Fig. S8c). Finally, in fermentable media, also nucleoid number was largely independent of mtDNA copy number (Fig. 3d). By contrast, in non-fermentable media, nucleoid number is higher in strains with higher mtDNA copy number (Fig. 3d). However, these changes are mild, and even in this condition, nucleoid number does not increase in direct proportion with mtDNA copy number. This suggests that nucleoid partitioning or organisation is buffered against changes in total mtDNA content.

**Figure 3:**
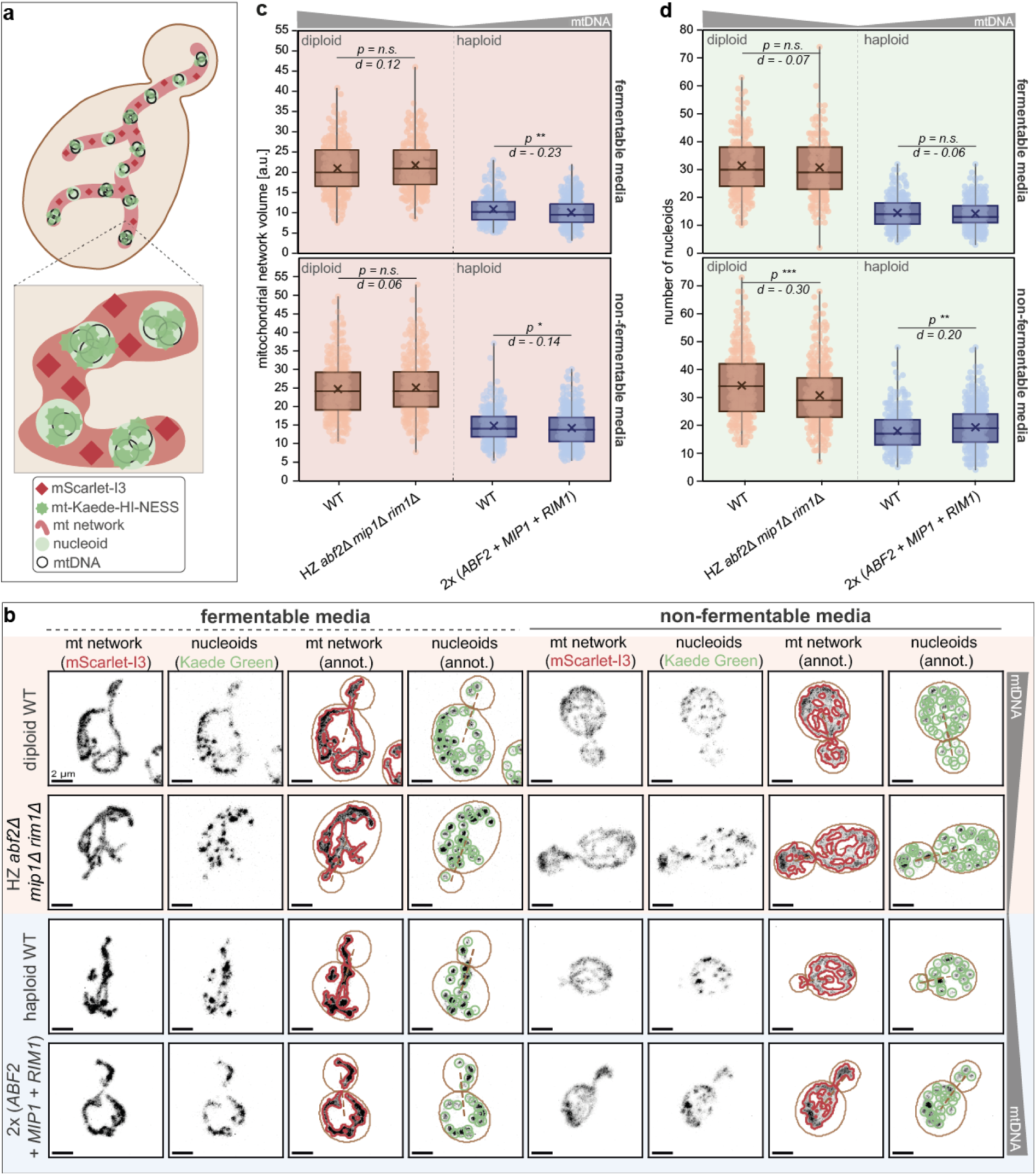
Mitochondrial network volume and nucleoid number are only weakly dependent on mtDNA copy number. **a)** The mt-Kaede-HI-NESS reporter is targeted to the mitochondrial matrix and binds mtDNA. The mitochondrial network was visualised with mScarlet-I3 targeted to the mitochondrial matrix. **b)** Representative confocal live-cell images (maximum projections) of the mitochondrial network (mScarlet-I3) and nucleoids (Kaede Green), as well as automatically obtained annotations of mitochondrial network (red), nucleoids (green) and cell segmentations (brown), for cells grown in fermentable (four left panels) and non-fermentable media (four right panels). Strains shown are diploid wild type (orange; top) and hemizygous deletion of *ABF2, MIP1,* and *RIM1* (orange; bottom); and haploid wild type (blue; top) and a strain with two copies of *ABF2, MIP1,* and *RIM1* (blue; bottom). Scale bars represent 2 µm. mtDNA concentrations in microscopy strains were confirmed by DNA-qPCR as shown in Fig. S8d. **c-d)** Analysis of mitochondrial network volume (**c**) and the number of nucleoids (**d**) from single cells microscopy in fermentable (top) and non-fermentable (bottom) media for diploid (orange) and haploid (blue) strains. Boxplots show the interquartile range (box), total range of data (whisker), median (line), and mean (cross). Number of single cells (dots) measured in three replicates: in fermentable media n_WT diploid_ = 313, n_HZ *abf2*Δ_*_mip1_*_Δ_*_rim1_*_Δ_ = 300, n_WT haploid_ = 247, n_2x (*ABF2*+*MIP1*+*RIM1*)_ = 528 (three individual clones pooled); in non-fermentable media: n_WT diploid_ = 333, n_HZ_ *_abf2_*_Δ*mip1*Δ*rim1*Δ_ = 331, n_WT haploid_ = 327, n*_2x (ABF2+MIP1+RIM1)_*= 581 (three individual clones pooled). Significance was determined compared to the respective wild type by a two-tailed t-test (* p<0.05, ** p<0.01, *** p<0.001), and effect sizes were calculated as Cohen’s d.

As an alternative approach, we used a well-established system based on fluorescently tagged LacI binding to multiple LacO repeats stably integrated into the mitochondrial genome^5,25^ and confirmed our finding that nucleoid number and network volume are independent of mtDNA copy number for cells grown in fermentable media (Fig S9).

### Expression of mtDNA-encoded transcripts increases with mtDNA copy number, and the corresponding proteins are partially dosage compensated

To better understand the cellular adaptation to altered mtDNA copy numbers, we asked how mtDNA copy number affects the expression of mtDNA-encoded genes. We analysed the transcriptomes of haploid and diploid strains with altered mtDNA copy number grown in non-fermentable media with RNA-sequencing, and the corresponding proteome with TMT mass spectrometry. To determine how expression of a given gene depends on mtDNA copy number, we sought to integrate the behaviour of all analysed haploid and diploid strains. To achieve this, we normalised transcript and protein abundances as well as mtDNA concentrations to the respective wild-type strains. For each gene, we then calculated the Spearman correlation coefficient of this relative transcript (or protein) abundance and the relative mtDNA concentration (Fig. 4a). While we find that for nuclear-encoded transcripts, this mtDNA-dependence is symmetrically distributed around 0 (mean -0.02), most mtDNA-encoded RNA and their mean show a positive correlation with mtDNA concentration (Fig. 4b-c) (mean 0.42). In addition to the eight mitochondrially encoded protein-coding open reading frames (ORFs), the W303-yeast mitochondrial genome encodes up to nine intron-derived proteins that function as maturases or endonucleases^48–50^. Although mitochondrially encoded proteins are difficult to detect by proteomics due to their hydrophobicity, we were able to detect six of these mitochondrially encoded proteins in our analysis. Interestingly, these proteins show a less clear dependence on mtDNA concentration (Fig. 4d; Fig. S10). Taken together, we found that altered mtDNA copy number leads to corresponding changes in the expression of mtDNA-encoded RNA, but that these changes in RNA levels are partially buffered on the protein level.

**Figure 4:**
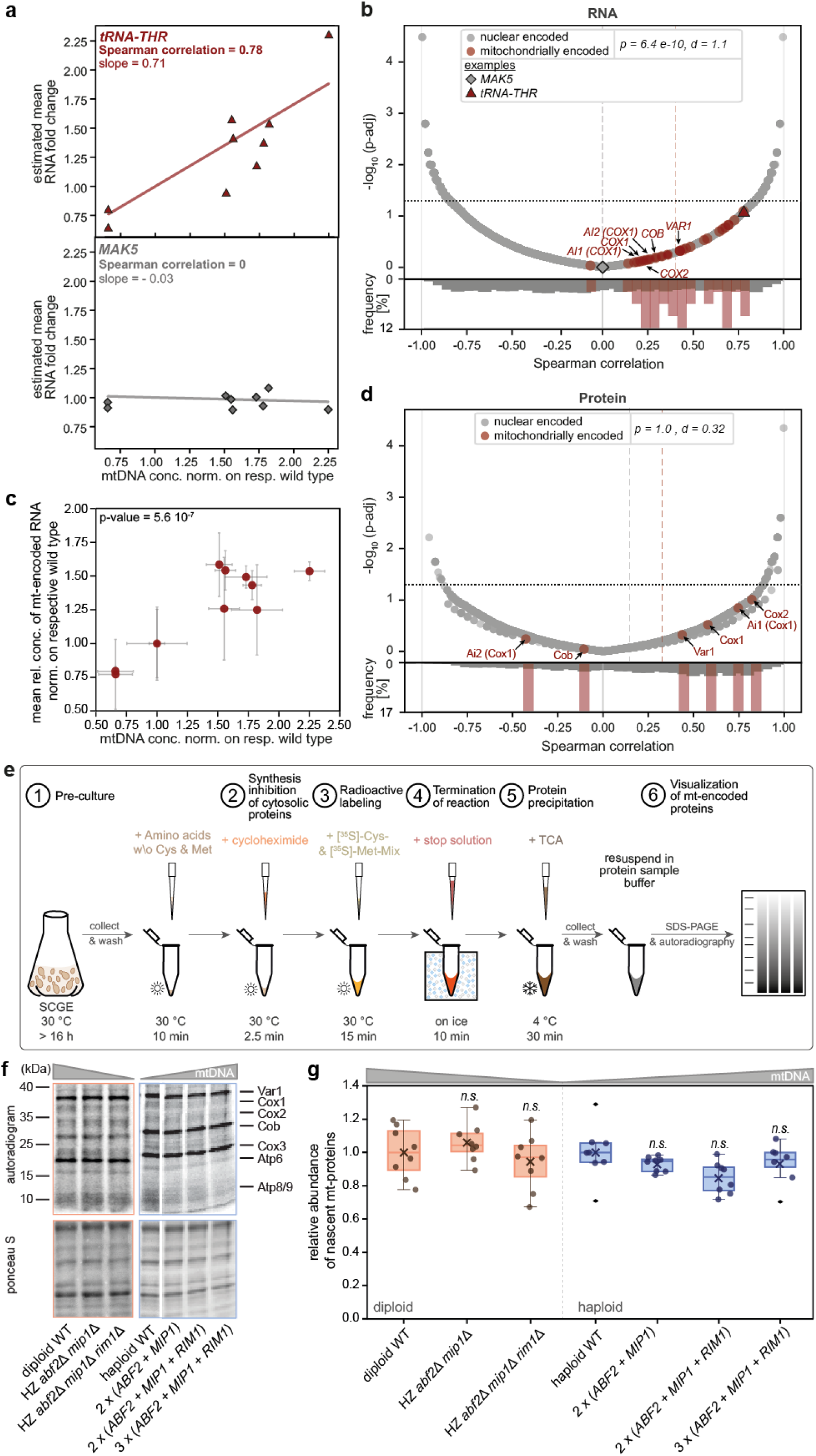
Higher mtDNA copy numbers lead to increased expression of mitochondrially encoded RNA, but the corresponding proteins are partially dosage compensated. **a)** Example of the estimated mean RNA fold change of at least three replicates in various strains as a function of the mtDNA concentration normalised on the respective wild type, for one mitochondrially encoded (tRNA-Thr, red; top) and one nuclear-encoded (MAK5, grey; bottom) RNA. **b)** Spearman correlation is used to characterise the dependence of the relative RNA concentration on mtDNA concentration for nuclear-encoded (grey) and mitochondrially encoded (red) genes. Volcano plot (top) shows negative logarithmic adjusted p-value as a function of the Spearman correlation, with the mean of each category shown as a dashed line. The threshold to significant p-values is shown as black dotted horizontal line. Representative examples of a) are highlighted. The distributions of Spearman correlation coefficients are shown as histograms (bottom). The significance between distributions was determined by a two-sided Fisher’s Exact test (* p<0.05, ** p<0.01, *** p<0.001), and effect sizes were calculated as Cohen’s d. **c)** Mean relative RNA-concentration of all mitochondrially encoded genes of at least three replicates as a function of the mtDNA concentration for haploid and diploid strains, both normalised on the respective wild type. Error bars indicate standard deviations. Significance was determined by Spearman correlation analysis. **d)** Spearman correlation is used to characterise the dependence of the relative protein concentration on mtDNA concentration for nuclear-encoded (grey) and mitochondrially encoded (red) proteins. Volcano plot (top) shows the negative logarithmic adjusted p-value as a function of the Spearman correlation, with the mean of each category shown as a dashed line. The threshold to significant p-values is shown as black dotted horizontal line. The distributions of Spearman correlation coefficients are shown as histograms (bottom), and mean Spearman correlation coefficients per category are shown as dashed lines. The significance between distributions was determined by a two-sided Fisher’s Exact test (* p<0.05, ** p<0.01, *** p<0.001), and effect sizes were calculated as Cohen’s d. **e)** Illustration of radiolabelling experiment: After preculturing, the cells are transferred into media including all amino acids except cysteine and methionine. Next, the synthesis of cytosolic proteins is inhibited with cycloheximide, followed by radioactive labelling of nascent mitochondrially encoded proteins by the addition of radioactively labelled cysteine and methionine. After the termination of the reaction by adding a stop solution, cells are precipitated, and mitochondrially encoded proteins are visualised by SDS-PAGE and autoradiography. **f)** Example of autoradiogram (top) and corresponding Ponceau S staining (bottom) for diploid (orange) and haploid (blue) strains. mtDNA-encoded proteins (right) are highlighted. **g)** Relative abundance of total nascent mtDNA-encoded proteins for diploid (orange) and haploid (blue) strains. Boxplots show the interquartile range (box), total range of data (whisker), median (line), and mean (cross). Single replicates are shown as dots. *S*tatistical significance was evaluated using one-way ANOVA with Tukey’s post hoc test, or, when assumptions were not met, a Kruskal–Wallis test followed by pairwise Wilcoxon tests with Benjamini–Hochberg correction.

To gain further insights into how mtDNA copy number affects the production of mtDNA-encoded proteins, we used radiolabelling experiments to monitor the abundance of nascent mtDNA-encoded proteins (Fig. 4e). The abundance of newly synthesised mtDNA-encoded proteins was largely unchanged across strains with different mtDNA copy numbers, suggesting that increases in mtDNA template availability do not proportionally increase mitochondrial translation output (Fig. 4f-g; Fig. S11-S12). This indicates that variations in mtDNA copy number are buffered by dosage compensation, most likely occurring at the translational level or by co-translational degradation.

### Expression of nuclear-encoded OXPHOS genes is coordinated with mtDNA copy number

Most mtDNA-encoded proteins are part of the oxidative phosphorylation complexes. To ensure the correct stoichiometry of the nuclear- and mtDNA-encoded subunits, cells coordinate their synthesis^51–53^. Since the mtDNA-encoded transcripts corresponding to subunits of the OXPHOS complexes increase with mtDNA copy number, we wondered how copy number changes affect the expression of nuclear-encoded OXPHOS genes. Consistent with a coordinated expression of these two sets of genes, we observed increased relative transcript concentrations of the nuclear-encoded subunits (Fig. 5a). Again, this increase with mtDNA copy number seems less pronounced on the protein level (Fig. 5b). Importantly, we did not observe a clear mtDNA dependence across all mitochondrial related genes, suggesting that this regulation is specific to the OXPHOS subunits (Fig. S13a-b).

**Figure 5:**
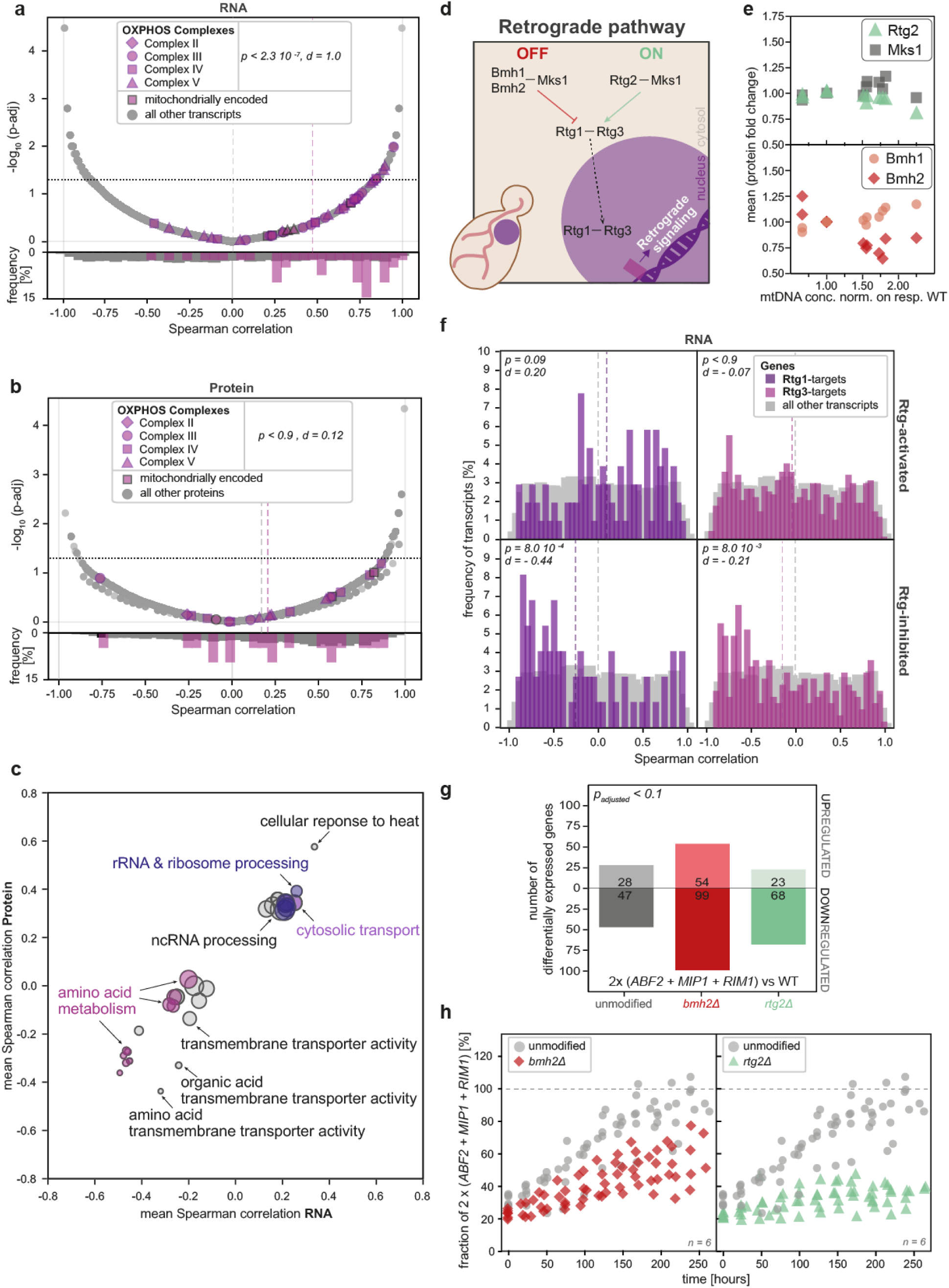
mtDNA copy number changes alter the expression of nuclear-encoded genes, partially through retrograde signalling. **a)** Spearman correlation is used to characterise the dependence of the relative RNA concentration on mtDNA concentration for genes encoding for subunits of the OXPHOS complexes (purple) as well as all other genes (grey). Mitochondrially encoded OXPHOS complex subunits are highlighted. Volcano plot (top) shows the negative logarithmic adjusted p-value as a function of the Spearman correlation. The threshold to significant p-values is shown as black dotted horizontal line. The distributions of Spearman correlation coefficients are shown as histograms (bottom), and the mean Spearman correlation coefficients per category are shown as dashed lines. The significance between distributions was determined by a two-sided Fisher’s Exact test (* p<0.05, ** p<0.01, *** p<0.001), and effect sizes were calculated as Cohen’s d. **b)** Spearman correlation is used to characterise the dependence of the relative protein concentration on mtDNA concentration for subunits of the OXPHOS complexes (purple) as well as all other proteins (grey). Mitochondrially encoded OXPHOS complex subunits are highlighted. Volcano plot (top) shows the negative logarithmic adjusted p-value as a function of the Spearman correlation. The threshold to significant p-values is shown as black dotted horizontal line. The distributions of the Spearman correlation coefficients are shown as histograms (bottom), and the mean Spearman correlation coefficients per category are shown as dashed lines. The significance between distributions was determined by a two-sided Fisher’s Exact test (* p<0.05, ** p<0.01, *** p<0.001), and effect sizes were calculated as Cohen’s d. **c)** Annotation enrichment analysis using the Spearman correlation coefficients characterising the dependence of the protein and relative RNA concentrations on the mtDNA concentration. Each dot shows an annotation group that is significantly enriched on both the protein and RNA levels and includes a minimum of 10 analysed proteins or RNAs (but not more than 500). The dot size represents the number of analysed proteins in each group, and the position is determined by the mean Spearman correlation of the group. **d)** Illustration of the retrograde pathway sensing mitochondrial dysfunction: If the pathway is switched off, Mks1 is highly phosphorylated and binds to Bmh1 and Bmh2, while if it is switched on, Mks1 is hypophosphorylated and binds to Rtg2. Additionally, highly phosphorylated Rtg3-proteins form complexes with Rtg1, and depending on the state of the pathway, this complex can translocate from the cytosol into the nucleus. Here, Rtg1/3 act as transcription factors, leading to a retrograde response. **e)** Mean protein fold change (two-replicates) as a function of the mtDNA concentration, both normalised on the respective wild type for Rtg2 (green triangle; top), Mks1 (grey square; top), Bmh1 (light red dot; bottom) and Bmh2 (red diamond; bottom). **f)** Histogram of the Spearman correlation coefficients for Rtg1- or Rtg3-dependent genes (purple and pink) and all other transcripts (grey) and, separated by genes that are activated (top) or inhibited (bottom). The mean Spearman correlation coefficients per category are shown as dashed lines. The significance between distributions was determined by a Kolmogorov-Smirnov test (* p<0.05, ** p<0.01, *** p<0.001), and effect sizes were calculated as Cohen’s d. Additional results on the protein level are shown in Figure S13c. **g)** Number of significantly differentially expressed genes comparing the strains with two copies of *ABF2, MIP1,* and *RIM1* to the corresponding haploid reference strains. Strains were otherwise unmodified (grey) or additionally carried deletions of *RTG2* (green) or *BMH2* (red). Categorised into up- or downregulated genes. **h)** Competition assays comparing growth of strains carrying two copies of *ABF2, MIP1,* and *RIM1* against the respective control strain for strain pairs that were otherwise unmodified (grey) or carried additional deletions of *RTG2* (green triangles; right) or *BMH2* (red diamonds; left). The two-copy strain initially constituted 25% of the mixture, and its dynamic development was monitored by DNA-qPCR. Symbols show individual measurements from six replicates.

Next, to identify potential other processes that are dependent on mtDNA copy number, we performed a genome-wide annotation enrichment analysis (Fig. 5c). Interestingly, this revealed that genes linked to rRNA and ribosome processing are upregulated in strains with higher mtDNA copy numbers, which may be linked to their increased growth rate. By contrast, genes related to amino acid metabolism are downregulated.

### Retrograde signalling is important for adaptation to increased mtDNA copy numbers

As our analysis revealed that some nuclear-encoded genes are regulated in an mtDNA copy number-dependent manner, we next asked how this regulation of nuclear gene expression is controlled. One possibility is the so-called retrograde pathway, which governs the nuclear localization and activity of the transcription factors Rtg1 and Rtg3 (Fig. 5d). To investigate whether retrograde signalling is also modulated by mtDNA copy number, we first analysed the relative concentrations of four key regulatory proteins of the retrograde pathway, Bmh1, Bmh2, Rtg2, and Mks1 (Fig. 5e). Interestingly, we found that the relative concentration of the retrograde pathway inhibitor Bhm2 decreases with mtDNA copy number, while the concentration of its paralog Bmh1 increases slightly. Guided by this observation, we then analysed the mtDNA-dependence of Rtg1- and Rtg3-target genes. Specifically, we investigated whether genes that were reported to be differentially expressed in either *RTG1* or *RTG3* deletion strains are expressed in an mtDNA-dependent manner. Indeed, we found that transcript levels of genes that are controlled by retrograde signalling are modulated by mtDNA copy number (Fig. 5f). Expression of genes reported to be inhibited by Rtg1 and Rtg3 is downregulated in cells with higher mtDNA copy number. Moreover, expression of genes activated by Rtg1 is upregulated, albeit not statistically significantly. This suggests that increased mtDNA copy numbers activate the retrograde pathway.

Since our findings suggest that mtDNA copy number changes affect retrograde signalling, we asked whether disrupting the pathway affects how cells adapt to mtDNA copy number changes. To test this, we individually deleted *BMH2* or *RTG2*, which are important regulators of the retrograde pathway, in the haploid wild type and the haploid strain with two copies of the three limiting factors. We did not observe any obvious effects of the two different deletions on mtDNA copy number, cell volume, or bud fractions (Fig. S13d-f). We then asked whether disruption of retrograde signalling alters the effect of mtDNA copy number changes on the transcriptome. RNA-sequencing analysis revealed that the disruption, in particular through the deletion of *BMH2*, leads to a higher number of differentially expressed genes in a strain with increased mtDNA copy number (Fig. 5g). This indicates that retrograde signalling may be important for the adaptation of cells to increased mtDNA copy numbers. To test whether intact retrograde signalling is required for the faster cell growth caused by increased mtDNA copy numbers, we again performed competition assays. Strikingly, disruption of retrograde signalling by deletion of either *RTG2* or *BMH2* diminished the growth advantage of cells with increased mtDNA copy number (Fig. 5h).

In summary, our results suggest that higher mtDNA copy numbers lead to a stronger activation of retrograde signalling. This signalling mediates adaptation to the copy number alterations, and is required for the growth advantage on non-fermentable media associated with increased mtDNA copy numbers (Fig. 6).

**Figure 6:**
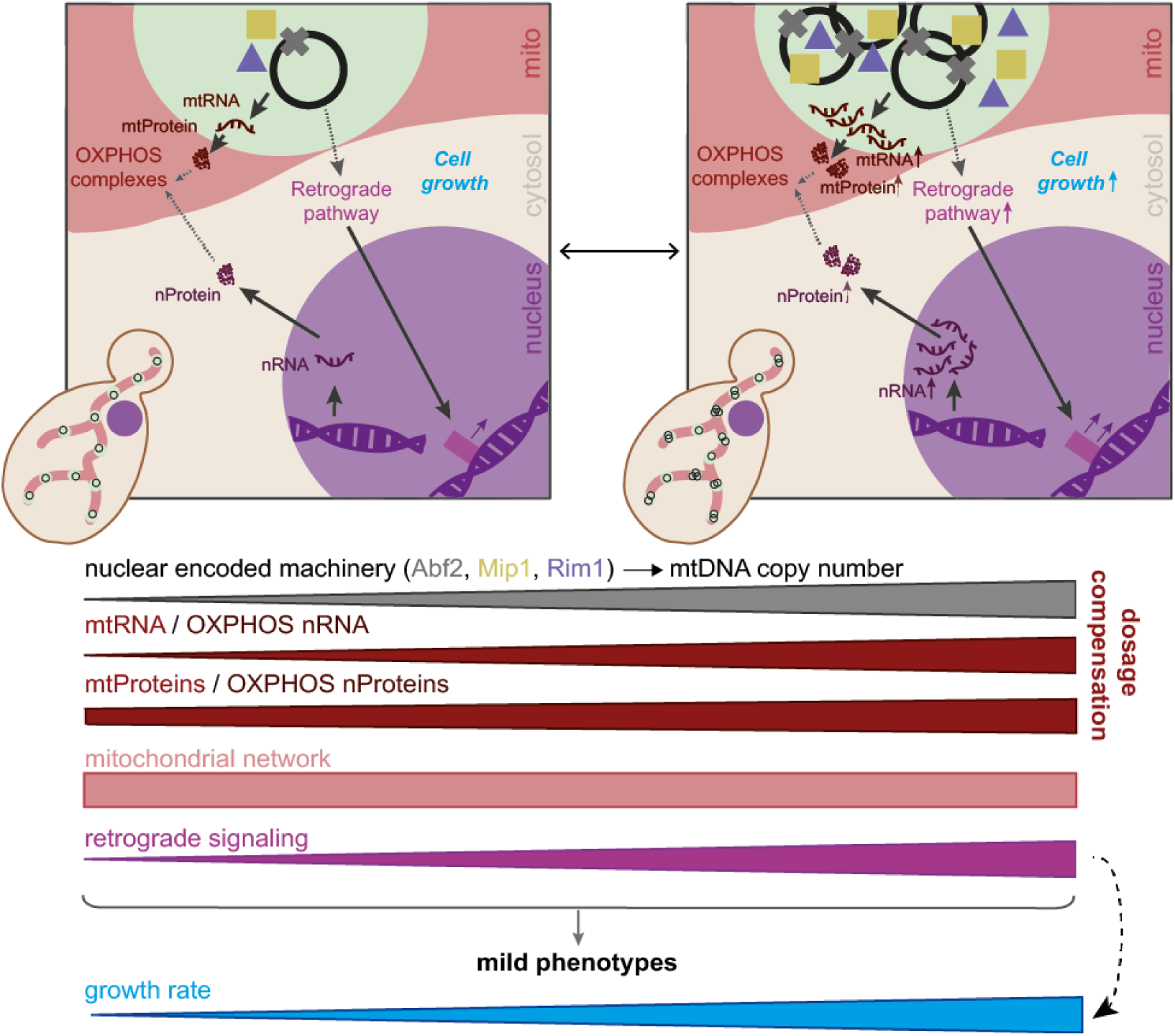
Illustration of the effect of mtDNA copy number changes on cell function. Cells are surprisingly robust to changes in mtDNA copy number induced by a manipulation of the limiting nuclear-encoded machinery (Abf2, Mip1, and Rim1). We attribute this to a combination of the dosage compensation of mtDNA-encoded proteins, the independent regulation of the mitochondrial network, as well as adaptation induced by retrograde signalling.

## Discussion

Through manipulation of gene copy numbers, we identified Abf2, Mip1, and Rim1 to be the three major nuclear-encoded factors that are limiting for mtDNA copy number in budding yeast. The amount of all three limiting proteins increases with cell volume, suggesting that a limiting machinery mechanism accounts for the coupling of mtDNA copy number to cell volume^5^. In this model, larger cells have more of the limiting proteins Abf2, Mip1, and Rim1, which in turn leads to a proportionally higher mtDNA copy number (Fig. 1g).

A similar limiting machinery mechanism may be operating in mammalian cells, where the homologs of Abf2, Mip1, and Rim1, namely TFAM, PolG, and mtSSB, have been reported to have a dosage-dependent effect on mtDNA copy number^54–60^. While this suggests that the limiting machinery mechanism we identified in budding yeast could be, in essence, conserved, it is important to note that the essential part of the limiting machinery mechanism is only the existence of limiting factors whose abundance increases with cell volume. Thus, the exact identity of the limiting factors may vary between organisms and even environmental conditions. For example, in mammals, the helicase Twinkle, which does not have a homolog in yeast, has also been reported to have a dosage-dependent effect on mtDNA copy number^61^.

Identification and genetic manipulation of the limiting machinery provided us with a powerful tool to investigate the consequences of mtDNA copy number changes on cell function. Surprisingly, we found that twofold up- or downregulation of mtDNA copy number only led to very mild phenotypic changes (Fig. 2). This suggests that while maintenance of mtDNA is critical, cells can buffer moderate copy number changes. Our observations are in line with a recent study that found that while decreased TFAM expression in mtDNA mutator mice leads to decreased mtDNA copy numbers, it only mildly affects their ageing phenotypes^62^.

One important factor that may be critical for the cellular robustness to mtDNA copy number changes is that mitochondrially encoded proteins are dosage compensated. While the transcript levels of mtDNA-encoded genes largely follow the mtDNA copy number (Fig. 4b-c), the relative concentration of the corresponding proteins is maintained more constant (Fig. 4d-g). This buffering at the protein level is consistent with the view^52,63^ that regulation of mtDNA-encoded proteins occurs mostly through translation and degradation rather than transcriptional regulation.

A second factor that may be important to maintain mitochondrial function despite moderate changes in mtDNA copy number is that mitochondrial morphology, in particular network volume, as well as the number of nucleoids, *i.e.,* the nucleoprotein complexes containing one or multiple mtDNA molecules, are largely independent of mtDNA copy number (Fig. 3). Supporting the observation that the mitochondrial network volume is independent of mtDNA copy number, we found that also the mitochondrial proteome mass fraction remains constant (Fig. S13c).

Consistent with the mild phenotypes we observed in strains with altered mtDNA copy number, we also found that the nuclear-encoded transcriptome and proteome exhibit only moderate changes (Fig. 5a-c). However, the fact that retrograde signalling is modulated as a function of mtDNA copy number suggests that cells sense and adapt to copy number changes (Fig. 5e-f). Accordingly, disrupting the regulation of retrograde signalling leads to a stronger modulation of the transcriptome in response to an mtDNA copy number increase (Fig. 5g). Critically, the faster growth we observed for cells with increased mtDNA copy number depends on intact retrograde signalling (Fig. 5h), indicating that this pathway is required for the cellular adaptation that enables the fitness benefit associated with higher mtDNA copy numbers.

Our finding that cells are robust to moderate changes in mtDNA copy number may be causally linked to the long-standing observation that cells, and even organisms, can tolerate mutations in a substantial fraction of their mtDNA copies^64^. For example, in humans, even large-scale deletions that lead to non-functional mtDNA copies only cause symptoms once they exceed a so-called heteroplasmy threshold of about 60-80%^65^. At the same time, this raises the question of where the threshold lies, at which increased or decreased mtDNA copy numbers begin to cause loss of robustness.

A main phenotype associated with mtDNA copy number changes we identified is that the growth rate in non-fermentable media increases with copy number (Fig. 2c-d). This feature is likely conserved in mammals, where faster tumour growth has been observed upon TFAM overexpression in xenograft tumour models developed from colorectal cancer cells^66^. Consistent with previous reports^31,32^, we found that higher mtDNA copy numbers also lead to reduced petite frequencies (Fig. 2e). At first glance, our findings therefore paradoxically suggest that increasing mtDNA copy number beyond wild-type levels increases cellular fitness. This raises the question of why the wild-type laboratory strain did not evolve to increase mtDNA copy number. Most likely, increased mtDNA copy number comes with a drawback that was not captured by our experiments. For example, purifying selection for functional mtDNA depends on the stochastic inheritance at cell division and is therefore faster at lower copy numbers^67,32^. High copy numbers may then lead to a problematic accumulation of detrimental mtDNA mutations.

In summary, we have introduced a novel model system to directly investigate the cellular consequences of mtDNA copy number alterations. We have shown that budding yeast can tolerate severalfold changes in mtDNA copy number, likely due to dosage compensation of mitochondrially encoded proteins as well as an mtDNA-independent regulation of network morphology and nucleoid number (Fig. 6). By mediating an adaptive response of nuclear gene expression, retrograde signalling plays an important role in the adaptation of cells to mtDNA copy number alterations. This retrograde response is also critical for the growth benefit in respiratory conditions we identified for cells with increased mtDNA copy number.

Given the robustness we observed here, it may seem surprising that mtDNA copy number changes are associated with many human diseases. This raises the question of whether it is not only the altered mtDNA copy number itself, but also a simultaneous failure of the adaptation response, which leads to pathological consequences. Moreover, it also remains to be investigated at what point the robustness of cells to mtDNA copy number changes fails, and which cellular processes are affected first.

## Methods

### Yeast strains

Yeast strains used in this study are listed in Supplementary Table 1. They are congenic with W303 and were constructed using standard methods. All manipulations were confirmed by PCRs and sequencing. Plasmids used to construct the strains are listed in Supplementary Table 2. All strains are available upon reasonable request.

### Yeast culturing

If not indicated differently, strains were grown at 30 °C at 250 rpm in a shaking incubator. All strains were inoculated in the appropriate media and cultured for 4 hours (synthetic complete media containing 2% glucose, SCD) or 6 hours (synthetic complete media containing 2% glycerol and 1% ethanol, SCGE), respectively. Then, cells were diluted to grow into the exponential phase until the next day, not exceeding OD_600_=1. Cells were then either harvested directly by centrifugation at 3,434 g for 5 minutes at 4 °C, or – where applicable – treated with appropriate concentrations of β-estradiol while growing in exponential phase overnight.

### mtDNA copy number measurements

To extract DNA, phenol-chloroform-isoamyl alcohol (PCI) extraction was performed as described by Seel et al.^5^. Briefly, the harvested cell pellet was resuspended in 200 µL DNA extraction buffer, pH 8.0 (2% Triton X-100, 1% SDS, 100 mM NaCl, 10 mM TRIS, 1 mM EDTA) and 200 µL PCI together with glass beads. Cells were mechanically disrupted by vortexing at 3,000 oscillations per minute. Next, the cells were centrifuged at 16,000 g for 5 minutes. The aqueous phase was then mixed with 500 µL 100% ethanol. After a repetition of the centrifugation, the cell pellet was washed using 700 µL 70% ethanol. Next, the cell pellet was dissolved in 50 µL nuclease-free water, mixed with 1 mg mL^-1^ RNase A (DNase-free) and incubated at 37 °C for 30 minutes to remove RNA residues. After adding 200 µL PCI and 200 µL DNA extraction buffer, the extraction steps were repeated. For qPCR, the DNA was diluted to 0.5 ng/µL after measuring the concentration via spectrophotometer at 260 nm. For amplification, a mastermix composed of a DNA-binding fluorescent dye (BioRad, SsoAdvanced Universal SYBR Green Supermix) and primer pairs for the nuclear DNA (nDNA) genes *ACT1* and *MRX6* and the mtDNA-encoded genes *COX2* and *COX3* were used (Supplementary Table 3). The qPCR protocol included an initial denaturation step of 10 minutes. All samples were analysed in technical triplicates, and downstream calculations were based on the mean Cq values of these replicates. Individual technical replicates were excluded when the corresponding standard deviation exceeded 0.5. Next, the gene-specific concentrations were quantified, and nDNA (*ACT1, MRX6*) as well as mtDNA (*COX2, COX3*) values were averaged to obtain pooled nDNA and mtDNA concentrations. mtDNA concentrations were normalised to nDNA to calculate the relative mtDNA copy number per nDNA unit. The budding index (percentage of budded cells) was determined by manual inspection and used to estimate the mean nDNA content per cell according to the following formulas:

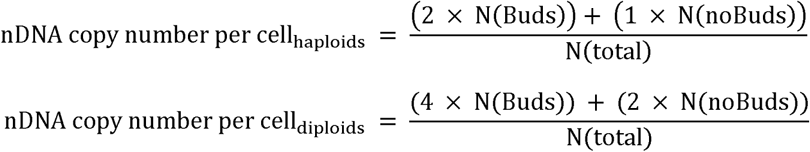

The mtDNA copy number per cell was then used to calculate the mtDNA concentration, by dividing by the cell volume, determined by Coulter counter measurements. Data distributions were tested for normality before statistical evaluation was performed using a two-handed two-tailed t-test with a significance threshold of α = 0.05.

### RNA measurements (RNA sequencing and RT-qPCR)

For RNA purification, cells were cultured as described above and 10 mL were harvested and stored on ice for less than 15 minutes until the RNA extraction was performed. For this, the YeaStar RNA Kit (Zymo Research) was used as specified in the instruction protocol. Afterwards, the RNA concentration was determined using a spectrophotometer, and the sample could be stored at -80°C until further analysis. To avoid DNA contamination, the samples were incubated with DNase I (Life Technologies) for 15 minutes. To stop the DNA digestion step, the samples were incubated with EDTA at 65 °C for 10 minutes.

Samples intended to be used for RNA sequencing were then rRNA depleted, cDNA transcribed, subjected to end repair and A-tailing, adaptor ligated and a library was amplified by Novogene Ltd. Library quality was assessed on an Agilent 5400 system (Agilent, USA and quantified by qPCR, followed by sequencing on a Illumina NovaSeq X Plus with a PE150 strategy. For RNAseq analysis, all genes detected across all independent sets of experiments were included. Counts were transcript per million (TPM)-normalised and log10-transformed after adding a pseudocount of 1 to avoid undefined values. Differential gene expression analysis was performed on raw counts using pyDESeq2^68^. Each strain was compared to its corresponding background strain. Significance was assessed using the Wald test, and differentially expressed genes were defined as those with an adjusted p-value < 0.1.

For RT-qPCR, for each sample 1 µg of RNA was transcribed to cDNA using the high-capacity cDNA reverse-transcription kit according to the protocol (Thermo Fisher Scientific). Next, the cDNA samples were diluted 1:10 for qPCR with specific primers, except for the ribosomal RNA *RDN18,* for which the samples were diluted 1:100. The RT-qPCR was performed using the fluorescent dye SYBR Green for detection. All samples were analysed in technical triplicates, and the resulting relative concentrations were derived by normalisation to *RDN18*.

### Measurements of cell volume, bud fraction, and doubling time

Mean cell volume of 20,000 to 50,000 cells in a range between 10 and 328 fL was measured using a Coulter counter, after sonicating the cells for 10 seconds.

Bud fraction, i.e. the percentage of cells with a bud, was determined by visual inspection using a microscope, after sonicating the cells for 10 seconds. A total number of at least 100 cells each was classified independently by two researchers and the results were pooled.

For analysis of the doubling time, cells were cultured as described above and diluted to a low concentration for the next day (OD_600_ < 0.1). A growth curve was obtained by measuring the optical density of the culture between 0.1 and 1 every 30 minutes. The doubling time was determined by calculating the slope of a linear regression fitted to the log-transformed data over time.

### Competition assays

To directly compare growth rates between two strains, competition assays were performed as follows: After culturing each strain for at least 24 hours as described above, each culture was diluted to an OD_600_ of 0.3 in 10 mL fresh media. Reference samples were obtained by harvesting 7 mL of these diluted cultures. Subsequently, each strain was mixed with its corresponding auxotrophy-corrected control strain at a 25:75 ratio. After mixing, a 7 mL aliquot was collected and stored for DNA extraction (T = 0 h). The remaining culture was diluted into fresh media and maintained in exponential growth at 30 °C and 250 rpm. For 5–11 consecutive days, cultures were adjusted daily to an OD_600_ of 0.3, after which 7 mL were harvested for analysis and the residual volume used to dilute the culture in fresh media. After DNA extraction of the samples using PSI-extraction and an RNase digestion step (see above), 1 ng was used for DNA-qPCR analysis, using the fluorescent dye SYBR Green and primer pairs for *ACT1* and *AMPR* (Supplementary Table 3). As the control strain does not include the *AMPR* sequence, the *AMPR* signal in the mixed culture reflects the fraction of the strain of interest. To determine this fraction, the mean Cq value of technical triplicates was used to calculate the ratio of *AMPR* and *ACT1* concentration. Values for each time point were normalised to the corresponding reference sample of the pure strain.

### Petite frequency assays

Strains were inoculated into 4 mL YPD and grown overnight at 30 °C and 250 rpm. The following day, 50 µL of each culture was transferred into 7 mL fresh YPD and incubated for an additional 2 hours. Optical density was measured using a NanoDrop One©, after which cells were washed and resuspended to an OD_600_ of 1 in 1 mL ddH_2_O. A total of 150 µL of this suspension was plated onto YPG media supplemented with 0.1% glucose. Two plates were prepared per strain and incubated at 30 °C for around 48 hours. Petite frequency was quantified by visually scoring at least 500 colonies per biological replicate.

### Stress spot assays

Before plating, the strains were inoculated in 4 mL YPD and grown overnight at 30 °C and 250 rpm. Optical density was measured using a NanoDrop One©, and cultures were adjusted to an OD_600_ of 1. After a washing step, the cells were transferred to 1 mL ddH_2_O and diluted in a five-step 1:10 series. Next, 5 µL of each dilution was spotted onto the appropriate agar plate. For temperature-sensitivity assays, YPD, SCD, or SCG plates were incubated at 23, 30, or 37 °C for ≥ 48 hours. For chemical stress assays, compounds were added to the media prior to plate casting: 0.6 M NaCl for salt stress on YPD plates, and 100 mM hydroxyurea for replicative stress on SCD plates. These plates were incubated at 30 °C until the wild-type control exhibited robust colony growth. Images were acquired using a transilluminator without UV illumination. Phenotypes were evaluated independently by three researchers under blinded conditions to ensure unbiased assessment.

### Time-lapse microscopy

After culturing the cells for at least 24 hours in the appropriate filtered media, they were diluted to OD_600_ = 0.1 in 5 mL. Time-lapse microscopy was carried out using a custom microfluidic device^40^ capable of immobilising single cells while continuously supplying fresh media. Growth media was supplied at 20 µL/min, and sample temperature was maintained at 30 °C using an objective and stage insertion heater. The analysis of steady-state growth was performed using non-fermentable SCGE media for the complete duration of the experiment. For the media-switch experiment, cells were cultured before, as well as for the first two hours of the microscopy experiment, in fermentable SCD media (2% glucose) (Fig. 2f). Afterwards, the media was switched to a minimal media containing low glucose concentrations SMM (0.05% glucose and 0.95% sorbitol), additionally supplemented with 1% tryptophan, 1% histidine, 1% leucine, 1% uracil. As the exchange of the media inside the microfluidic chamber takes approximately 30 minutes, the time of media switch was set to 2.5 hours of the experiment for downstream analysis. Imaging was performed on a Nikon Eclipse Ti-E microscope (NIS-Elements) equipped with an Andor iXon Ultra 888 camera and SPECTRA X light engine illumination. Phase-contrast images were acquired using a Plan Apo λ 100x/1.45 NA Ph3 oil-immersion objective with an additional 1.5x magnifier, a 100 millisecond exposure time and a 3-minute imaging interval to generate time-lapse sequences. For each strain and biological replicate, five positions were measured in parallel, filled with not more than six cells in frame 1, to avoid overcrowding. Abnormally shaped buds were determined for S/G2/M-switchers with a bud size smaller than 26 fL when facing the media switch.

### Live-cell confocal microscopy for analysis of mitochondrial network and nucleoids

Live-cell confocal microscopy was used to quantify mitochondrial network morphology and nucleoid number. The chambers of a µ-Slide 8 Well ibi-Treat were coated with Concanavalin A (ConA; 1 mg/mL in H_2_O) by adding 200 µL per well and incubating for 10 minutes at room temperature. Slides were washed twice, air-dried, and prepared for cell loading. Cultured cells (see above) were sonicated for 10 seconds, then washed twice and transferred into fresh filtered media. Then, 200 µL of the cells were added to each well and incubated at room temperature for 5 minutes to allow for cell adhesion. Next, the supernatants were removed, wells were washed twice, and 200 µL of fresh media was added. Imaging was performed on a Zeiss LSM 800 confocal microscope (Zen 2.3, Blue Edition).

Two different strategies were used to visualise the mitochondrial network and nucleoids. The first one visualises the mitochondrial network Su9-mt-Kaede-HI-NESS expressed from an *ACT1* promoter and *CYC1* terminator (excitation 488 nm; detection 410-546nm) and the mitochondrial network using mScarlet-i3 targeted to the mitochondrial matrix (excitation 561 nm; detection 583-700 nm)^47,69^. Images were acquired in confocal mode using a 63x/1.4 NA oil-immersion DIC objective with 0.23 µm z-steps over a 12 µm depth (haploids) or 15 µm depth (diploids). Bright-field images were obtained using the transmitted-light detector (T-PMT).

The second approach uses mKate2 targeted to the mitochondrial matrix as a fluorescent reporter for the mitochondrial network visualisation (excitation 561 nm; detection 510–700 nm), and LacI-2xmNeon binding to LacO repeats integrated into the mitochondrial genome for the nucleoids (excitation 488 nm; detection 410–546 nm)^5,25^. Images were acquired in confocal mode using a 63x/1.4 NA oil-immersion DIC objective with 0.35 µm z-steps over a 15.05 µm depth. Bright-field images were obtained using the transmitted-light detector (T-PMT).

### Analysis of microscopy data

All microscopy data were evaluated using the open-source software Cell-ACDC^41^ and SpotMAX^70^. Raw image sequences were first aligned and cropped to define the region of interest. Cells were then automatically segmented and tracked in the phase-contrast channel using YeaZ^71^ or cellpose v4^72^, embedded in Cell-ACDC. Segmentation and tracking outputs were manually curated, including assignment of buds to their corresponding mother cells and annotation of cell-division events.

The time-lapse microscopy datasets were exported for downstream analysis, which was performed primarily using code from Chadha et al.^43^. For the cells facing the media switch, buds with abnormal morphology were identified by visual inspection. Quantification of mitochondrial network morphology and nucleoid number from confocal microscopy was carried out using SpotMAX^70^ integrated within the Cell-ACDC software. The analysis parameters of the SpotMAX analysis are available at https://hmgubox2.helmholtz-muenchen.de/index.php/s/94Z84tXkaQxsBek. Note that due to the optical resolution limit, the width of the mitochondrial network could not be measured. The segmented voxels are therefore only a proxy (in arbitrary units) for the mitochondrial network volume^5,7^.

### Measurement of Proteome by TMT mass spectrometry

Peptide preparation was performed as described by Lanz et al.^28^. In brief, cell pellets were first lysed in Tris-HCl buffer (50 mM, pH 8.0) containing 0.2% Tergitol, 150 mM NaCl, 5 mM EDTA, and protease inhibitors using glass bead disruption. Protein concentration was determined by BCA assay. Proteins were reduced with DTT, alkylated with iodoacetamide, and precipitated using acetone/ethanol/acetic acid. Precipitated proteins were resuspended in 8 M urea buffer and digested with trypsin (1:50, overnight, 37 °C). Peptides were acidified with TFA, clarified by centrifugation, desalted using C18 columns, eluted with 80% acetonitrile, dried, and resuspended in water.

For TMT labelling, peptide samples were resuspended in 100 mM TEAB. 20 μg of peptide was labelled using 100 μg of TMTpro 16plex™ in a reaction volume of 25 μl for 1 hour. The labelling reaction was quenched with 3 μL of 5% hydroxylamine for 15 minutes. Labelled peptides were pooled, acidified to a pH of ∼2 using drops of 10 % trifluoroacetic acid, and desalted with a C18 column as described above. TMT-labelled peptides were fractionated using the Pierce High pH Reversed-Phase Peptide Fractionation kit. After fractionation, dried peptides were reconstituted in 0.1 %TFA. Peptide concentrations were determined using a Nanodrop before mass spectrometry analysis.

Desalted TMT-labelled peptides were analysed on a Fusion Lumos mass spectrometer (Thermo Fisher Scientific, San Jose, CA) equipped with a Thermo EASY-nLC 1200 LC system (Thermo Fisher Scientific, San Jose, CA). Peptides were separated by capillary reverse phase chromatography on a 25 cm column (75 μm inner diameter, packed with 1.6 μm C18 resin, AUR2-25075C18A, Ionopticks, Victoria Australia). Electrospray Ionization voltage was set to 1550 volts. Peptides were introduced into the Fusion Lumos mass spectrometer using a two-step linear gradient with 6–33% buffer B (0.1% (v/v) formic acid in 80% acetonitrile) for 145 minutes, followed by 33-45% buffer B for 15 minutes at a flow rate of 300 nL/min. Column temperature was maintained at 40 °C throughout the procedure. Xcalibur software (Thermo Fisher Scientific) was used for data acquisition, and the instrument was operated in data-dependent mode. Survey scans were acquired in the Orbitrap mass analyser over the range of 380 to 1800 m/z with a mass resolution of 70,000 (at m/z 200). Ions with a charge state of either 2, 3 or 4 were selected for fragmentation within an isolation window of 0.7 m/z. Selected ions were fragmented by Higher-energy Collision-induced dissociation (CID) with normalised collision energies of 35%, and the tandem mass spectra was acquired in the Ion trap mass analyser with a “Rapid” scan rate. Repeated sequencing of peptides was kept to a minimum by dynamic exclusion of the sequenced peptides for 60 seconds. For MS/MS, the AGC target was set to “Standard” and max injection time was set to 35 ms. Relative changes in peptide concentration were determined at the MS3-level by isolating and fragmenting the 95 most dominant MS2 ion peaks using HCD. TMT reporter ions were resolved in the Orbitrap at a resolution of 50,000.

All raw files were searched using MaxQuant (v21.6.7.0). A Reporter ion MS3 search was conducted using 16plex TMT isobaric labels. Variable modifications included oxidation (M) and protein N-terminal acetylation. Carbamidomethyl (C) was a fixed modification. The number of modifications per peptide was capped at five. Digestion was set to tryptic (proline-blocked). Database search was conducted using the UniProt proteome - Yeast_UP000002311_559292.

### Analysis of mtDNA copy number-dependence based on RNA sequencing and mass spectrometry experiments

To investigate the dependence of gene expression on mtDNA copy number (Figs. 4 & 5), the fold change values were obtained from the log2fold change (LFC) values received from *pyDESeq2*^68^. To clarify, the fold change values represent estimates from a negative binomial generalised linear model with replicate as a covariate. For the analogous analysis of mtDNA-dependence on the protein level, the fold change was defined as the ratio of the mean protein abundance between the strain and its corresponding wild type across all replicates. To understand whether the fold change for a given RNA or protein increases or decreases monotonically with the mtDNA copy number, we calculated the corresponding Spearman correlation coefficient as well as the associated p-value with the *Spearman* function from *scipy.stats*^73^. To correct for multiple testing, the adjusted p-values are calculated using the *multipletests* function from *statsmodels.stats.multitest*^74^, using the Benjamini/Hochberg (*fdr_bh*) parameter. For this analysis, all genetically manipulated genes were removed. For statistical analysis, a two-sided Fisher’s Exact Test was performed using *fisher_exact* from *scipy.stats*^73^. Additionally, to test whether the distributions of gene or protein subsets differed significantly from the remaining genes or proteins, a two-sample Kolmogorov-Smirnov test was applied using *ks_2samp* from *scipy.stats*^73^ with default settings. These results were not shown in the figures, as the significances were comparable to the p-values of the two-sided Fisher’s Exact Test. Targets of Rtg1 and Rtg3 used in Fig. 5f originated from Yeastract^75^, while mitochondrial-related genes were obtained from the Saccharomyces Genome Database^76^. Note that the manipulated limiting factors, genes corresponding to auxotrophic markers used for deletion or integration of additional copies of the limiting factors, as well as *TVP18*, part of which is encoded by the *ABF2* promoter sequence, were excluded from genome-wide analyses.

### Annotation Enrichment Analysis

Annotation enrichment analysis for proteins and RNAs with regard to the Spearman correlation coefficient characterising their mtDNA dependence was performed as described before^28,77^. Protein and RNA annotation groups were considered as significantly enriched if the Benjamin-Hochberg FDR was lower than 0.02. Enriched annotation groups were only considered if they contained at least 10 measured proteins or RNAs, but not more than 500.

### Proteome Mass Fraction Analysis

To calculate the proteome mass fraction of each mapped protein, we utilised the peptide feature information in MaxQuant’s evidence.txt output file. Each row of the evidence.txt file represents an independent peptide and its corresponding MS3 reporter ion measurements. Peptides without a signal in any of the TMT channels were excluded. Peptide measurements were assigned to a protein based on MaxQuant’s “Leading razor protein” designation. For each individual peptide measurement (i.e., each row in the evidence table), the fraction of ion intensity in each TMT channel was calculated by dividing the “Reporter ion intensity” column by the sum of all reporter ion intensities. To correct for loading differences between the TMT channels, each reporter ion channel was then normalised by dividing the fraction of ion intensity in each channel by the median fraction for all measured peptides (i.e., the median value for each column). This normalisation scheme ensures that each individual peptide measurement is equally weighted when correcting for loading error. To calculate proteome mass fractions, the MS1 precursor ion intensity of each peptide measured (the “Intensity” column in the evidence.txt table) was distributed between the individual MS3 reportion channels according to the loading-normalised value described. Protein-level ion intensities were then calculated for each TMT channel by summing together all peptides ion intensities for each protein. To estimate proteome mass fraction of the mitochondria, the protein-level ion intensities for annotated mitochondrial proteins were summed together and divided by the summed ion intensity of the entire proteome. Proteins included into the mitochondrial proteome mass fraction are those that are reported to be exclusively located in mitochondria.

### ^35^S methionine radiolabelling of nascent mitochondrially encoded proteins

Radiolabelling of nascent mitochondrially encoded proteins was performed according to previously described procedures^78^, including minor adaptations. Overnight cultures were grown in SCGE media for 16 hours and subsequently used to inoculate main cultures in SCGE to an OD_600_ of 0.1. Cultures were incubated at 30 °C with 140 rpm in a 50 mm orbit shaking incubator and harvested at mid-exponential phase (OD_600_≈ 1.5) by centrifugation at 4,000 g for 5 minutes at room temperature (RT). Cells were washed three times with ultrapure water, followed by two washes with SCGE media lacking amino acids. The OD_600_ was measured, and cell material corresponding to an OD_600_ of 4 was collected by centrifugation (10,000 g, 2 min, RT). Pellets were resuspended in 1.5 ml SCGE media without amino acids. Amino acids were supplemented to a final concentration of 12 μg/ml for all amino acids, except cysteine (7.5 μg/ml), while methionine was omitted. Cultures were incubated for 10 minutes at 30 °C with shaking at 600 rpm. Cycloheximide was then added to a final concentration of 0.5 mM, and incubation was continued for 2.5 minutes. Radiolabelling of nascent mitochondrially encoded proteins was induced by adding 3 μl of protein labelling mix (11 mCi/ml; [^35^S]-L-methionine and [^35^S]-L-cysteine, EasyTag™Express, Revvity;NEG772002MC). To ensure an exact labelling period of 15 minutes, the [^35^S] mix was added with a 10 second offset across samples, and harvesting was performed with the same offset. For termination, 400 μl of each sample was mixed with 100 μl of stop solution (1.5 M NaOH, 1 M β-mercaptoethanol, 20 mM PMSF, 30 mM non-radioactive L-methionine) and placed on ice for 10 minutes. For protein precipitation, trichloroacetic acid (TCA) was added to a final concentration of 14 % and samples were incubated for 30 minutes at 4 °C. Precipitates were collected by centrifugation (20,000 g, 20 min, 4 °C), washed with 1 ml 100 % acetone, and centrifuged again (20,000 g, 20 min, 4 °C). Pellets were resuspended in 150 μl protein sample buffer (4% SDS, 100 mM Tris, 20% glycerol, 100mM DTT, pH 6.8) and analysed by SDS-PAGE. Proteins were transferred to nitrocellulose membranes, and loading was verified by Ponceau S staining. Membranes were dried and exposed for autoradiography using an Amersham Typhoon scanner. Bands corresponding to individual mitochondrially encoded proteins were quantified with ImageJ and normalised to Ponceau S signals. Protein abundance (total or individual) is reported as fold change relative to wild-type samples. Box plots indicate mean (square), median (line), lower and upper quartiles, and whiskers representing the maxima and minima with the 1.5x interquartile range (IQR); individual data points are shown, and outliers (outside 1.5x IQR) are marked as red diamonds. Normality was assessed using the Shapiro–Wilk test, and homogeneity of variances with Levene’s test. Statistical significance was evaluated using one-way ANOVA with Tukey’s post hoc test, or, when assumptions were not met, a Kruskal–Wallis test followed by pairwise Wilcoxon tests with Benjamini–Hochberg correction.

## Acknowledgments

We thank Christof Osman, Dejana Mokranjac and the members of the Institute of Functional Epigenetics for valuable discussions. We thank Vineesh Suresh for initial exploratory and statistical analyses of the RNA-seq data, and Martin Wenninger for initial exploratory strain constructions and experiments. This work was funded by the Deutsche Forschungsgemeinschaft (DFG, German Research Foundation) through projects 416098229 and 431480687, by the Helmholtz Gemeinschaft, and by the European Union (ERC, MITOSIZE, 101169998). Views and opinions expressed are however those of the authors only and do not necessarily reflect those of the European Union or the European Research Council. Neither the European Union nor the granting authority can be held responsible for them. This work was supported by the Initiative and Networking Fund of the Helmholtz Association under the call HIRS, the EpiCrossBorders: International Helmholtz-Edinburgh Research School for Epigenetics. A.K. acknowledges funding from Kempestiftelserna (JCSMK23-0157), Konung Gustaf V:s och Drottning Victorias Stiftelse, the Swedish Research Council / Vetenskapsrådet (2025-05122) and the Strategic Research Grant for Scientifically Young Researchers from the Faculty of Medicine, Umeå University.

## Notes

### Competing Interest Statement

The authors have declared no competing interest.

