## Supplementary Information for "Retrograde signalling mediates cellular adaptation to mitochondrial DNA copy number alterations"

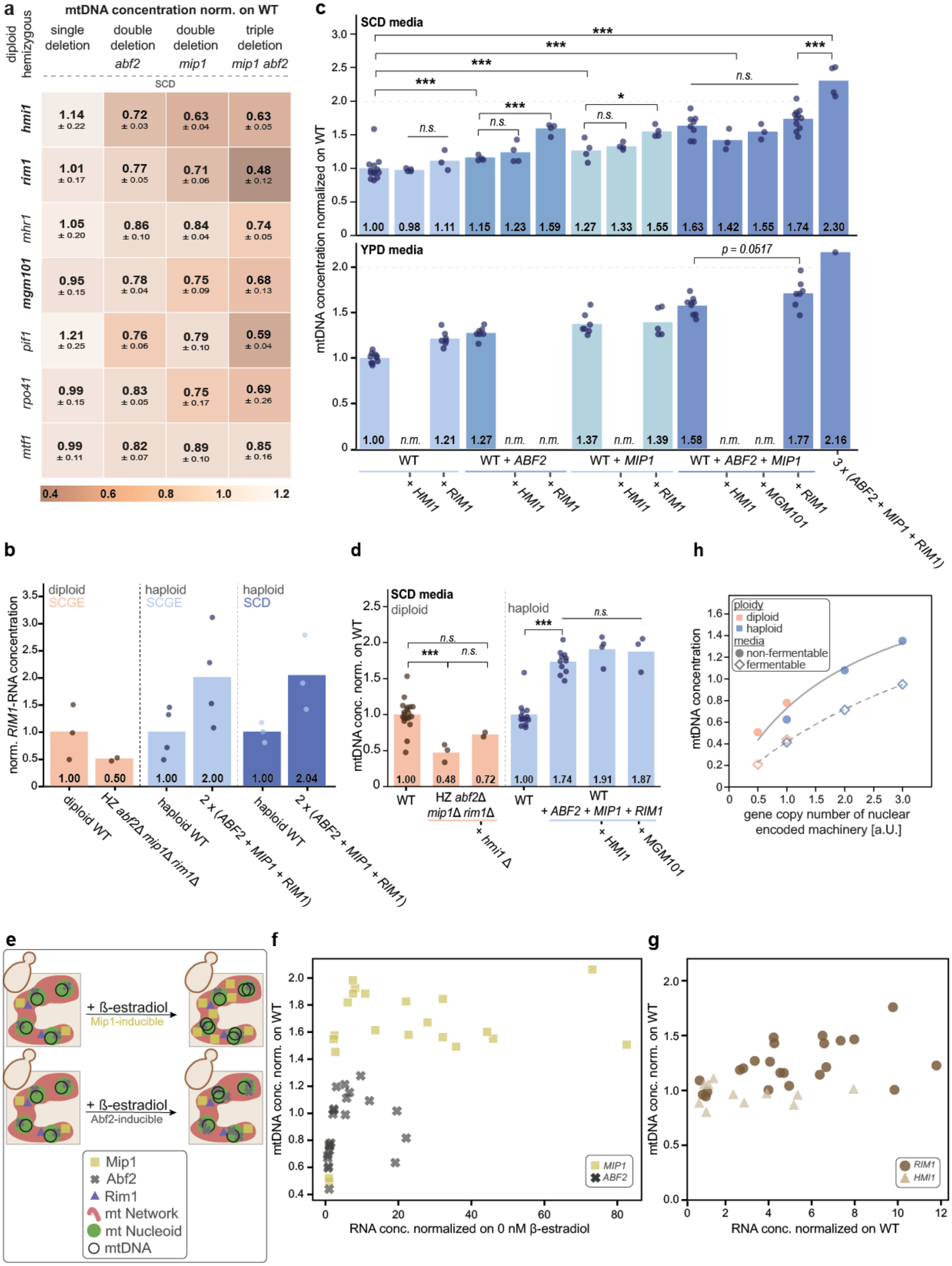
a
mtDNA concentration norm. on WT
diploid hemizygous
single deletion
double deletion abf2
double deletion mip1
triple deletion mip1 abf2
SCD
hmi1 1.14 ± 0.22 0.72 ± 0.03 0.63 ± 0.04 0.63 ± 0.05
rim1 1.01 ± 0.17 0.77 ± 0.05 0.71 ± 0.06 0.48 ± 0.12
mhr1 1.05 ± 0.20 0.86 ± 0.10 0.84 ± 0.04 0.74 ± 0.05
mgm101 0.95 ± 0.15 0.78 ± 0.04 0.75 ± 0.09 0.68 ± 0.13
pif1 1.21 ± 0.25 0.76 ± 0.06 0.79 ± 0.10 0.59 ± 0.04
rpo41 0.99 ± 0.15 0.83 ± 0.05 0.75 ± 0.17 0.69 ± 0.26
mtf1 0.99 ± 0.11 0.82 ± 0.07 0.89 ± 0.10 0.85 ± 0.16
0.4 0.6 0.8 1.0 1.2
c
SCD media
YPD media
mtDNA concentration normalized on WT
***
n.s.
*
p = 0.0517
1.00 0.98 1.11 1.15 1.23 1.59 1.27 1.33 1.55 1.63 1.42 1.55 1.74 2.30
1.00 n.m. 1.21 1.27 n.m. n.m. 1.37 n.m. 1.39 1.58 n.m. n.m. 1.77 2.16
WT
WT + ABF2
WT + MIP1
WT + ABF2 + MIP1
+ HMI1
+ RIM1
+ MGM101
3 x (ABF2 + MIP1 + RIM1)
b
norm. RIM1-RNA concentration
diploid SCGE
haploid SCGE
haploid SCD
1.00 0.50 1.00 2.00 1.00 2.04
diploid WT
HZ abf2Δ mip1Δ rim1Δ
haploid WT
2 x (ABF2 + MIP1 + RIM1)
haploid WT
2 x (ABF2 + MIP1 + RIM1)
d
SCD media
diploid
haploid
mtDNA conc. norm. on WT
n.s.
***
1.00 0.48 0.72 1.00 1.74 1.91 1.87
WT
HZ abf2Δ mip1Δ rim1Δ
+ hmi1 Δ
WT
WT + ABF2 + MIP1 + RIM1
+ HMI1
+ MGM101
h
ploidy
diploid
haploid
media
non-fermentable
fermentable
mtDNA concentration
gene copy number of nuclear encoded machinery [a.U.]
e
+ ß-estradiol
Mip1-inducible
+ ß-estradiol
Abf2-inducible
Mip1
Abf2
Rim1
mt Network
mt Nucleoid
mtDNA
f
mtDNA conc. norm. on WT
RNA conc. normalized on 0 nM β-estradiol
MIP1
ABF2
g
mtDNA conc. norm. on WT
RNA conc. normalized on WT
RIM1
HMI1

**Supplementary Figure S1: mtDNA copy number is set by nuclear-encoded machinery (Abf2, Mip1, Rim1) in fermentable media.**

**a)** mtDNA concentration normalised on diploid wild type (WT) for diploid hemizygous deletion strains determined by DNA-qPCR. Strains grown in fermentable media (SCD). Shown are means of at least three replicates per strain and standard deviations.

**b)** *RIM1*-RNA concentrations normalised on respective WT for strains grown in fermentable (SCD; dark blue) and non-fermentable media (SCGE; orange and light blue), determined by RT-qPCR. Shown are single replicates (dots) and the mean of all replicates (bars).

**c)** mtDNA concentration normalised on haploid WT for strains with additional copies of one or multiple genes grown in two different fermentable media (SCD on top; YPD on bottom), determined by DNA-qPCR. Shown are single replicates (dots) and the mean of all replicates (bars). Significance was determined by a two-tailed t-test (* p<0.05, ** p<0.01, *** p<0.001). Strains that were not measured are indicated as "not measured" (n.m.).

**d)** mtDNA concentration normalised on respective WT for strains grown in fermentable media (SCD) determined by DNA-qPCR. Shown are single replicates (dots) and the mean of all replicates (bar) for diploids (orange) and haploids (blue). Significance was determined by a two-tailed t-test (* p<0.05, ** p<0.01, *** p<0.001).

**e)** Illustration of the Mip1 or Abf2 strains overexpression system: A β-estradiol-inducible promotor is integrated in front of the endogenous *MIP1* (or *ABF2*) Open reading frame. Addition of higher concentrations of β-estradiol to the culture media leads to higher expression of *MIP1* (*ABF2*). The expression level was determined on RNA-level by RT-qPCR, and the resulting changes in mtDNA copy numbers were measured by DNA-qPCR. Results are shown in (**f**).

**f)** mtDNA concentrations normalised on WT increase with increasing RNA concentrations of *ABF2* (grey crosses) and *MIP1* (yellow squares). Increasing RNA concentrations were obtained in strains expressing *ABF2* or *MIP1* from an inducible promoter by increasing β-estradiol concentrations and normalised on 0 nM β-estradiol. At least two replicates per strain were performed in non-fermentable media (SCGE), and both mtDNA and RNA concentrations were measured by qPCR.

**g)** mtDNA concentrations normalised on WT increase with increasing RNA concentrations of *RIM1* (brown dot) and *HMI1* (light brown triangle). Increasing RNA concentrations were obtained by multicopy strains harbouring multiple copies of the respective gene. At least two replicates per strain were performed in non-fermentable media (SCGE), and both mtDNA and RNA concentrations were measured by qPCR.

**h)** mtDNA concentration increases with increasing gene copy number of the nuclear-encoded machinery (*ABF2*, *MIP1*, *RIM1*) in both fermentable (dots) and non-fermentable media (diamonds). The lines show Michaelis-Menten-like fits for fermentable media (dashed line; maximum of 2.6; Michaelis constant of 5.2) and non-fermentable media (straight line; maximum of 2.3; Michaelis constant of 2.1).

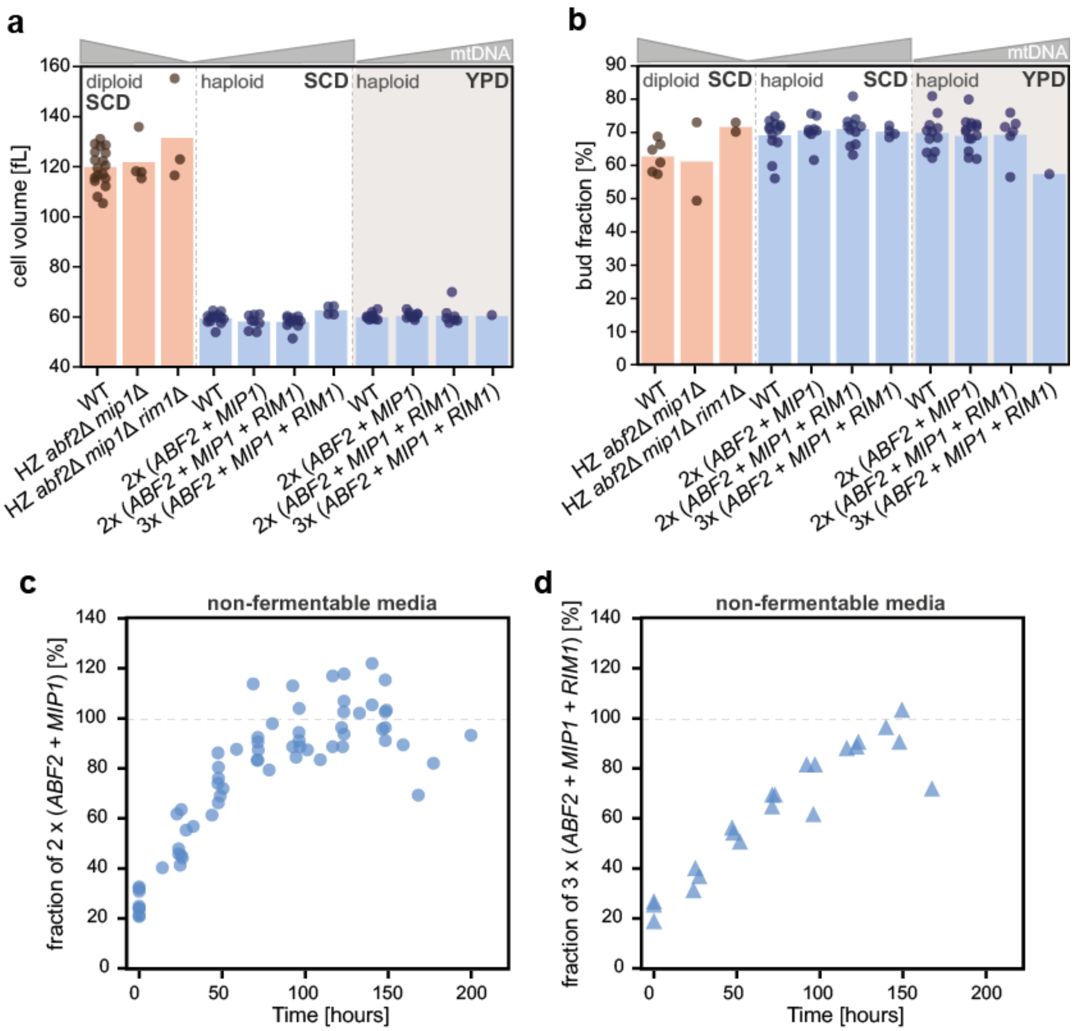


**Supplementary Figure S2: Cells are robust to mtDNA copy number changes in fermentable media and grow faster in non-fermentable media when they have higher mtDNA copy numbers.**

**a-b)** Cell volume and bud fraction measurements for various strains grown in fermentable media (SCD white background; YPD brown background). Diploids (orange) include wild type (WT), a hemizygous deletion of *ABF2* and *MIP1*, as well as a hemizygous deletion of *ABF2, MIP1,* and *RIM1*; haploids (blue) include WT, a strain with two copies of *ABF2* and *MIP1*, as well as strains with two or three copies of *ABF2, MIP1,* and *RIM1*. Three independent clones were measured and pooled for the strains with two copies of the three factors. Shown are single replicates (dots) and the mean of all replicates (bars). **a)** Mean cell volume determined by Coulter counter; **b)** Percentage of bud fraction determined by visual inspection under the microscope.

**c-d)** Competition assays testing the growth in non-fermentable media of strains against respective auxotrophy-corrected control strains. The fraction of the analysed strain in the mixture was initially 25%, and its dynamic development was monitored by DNA-qPCR. Spots show individual measurements from three replicates. **c)** Growth of a strain with two copies of *ABF2* and *MIP1* (three individual clones pooled) **d)** Growth of a strain with three copies of *ABF2, MIP1,* and *RIM1*.

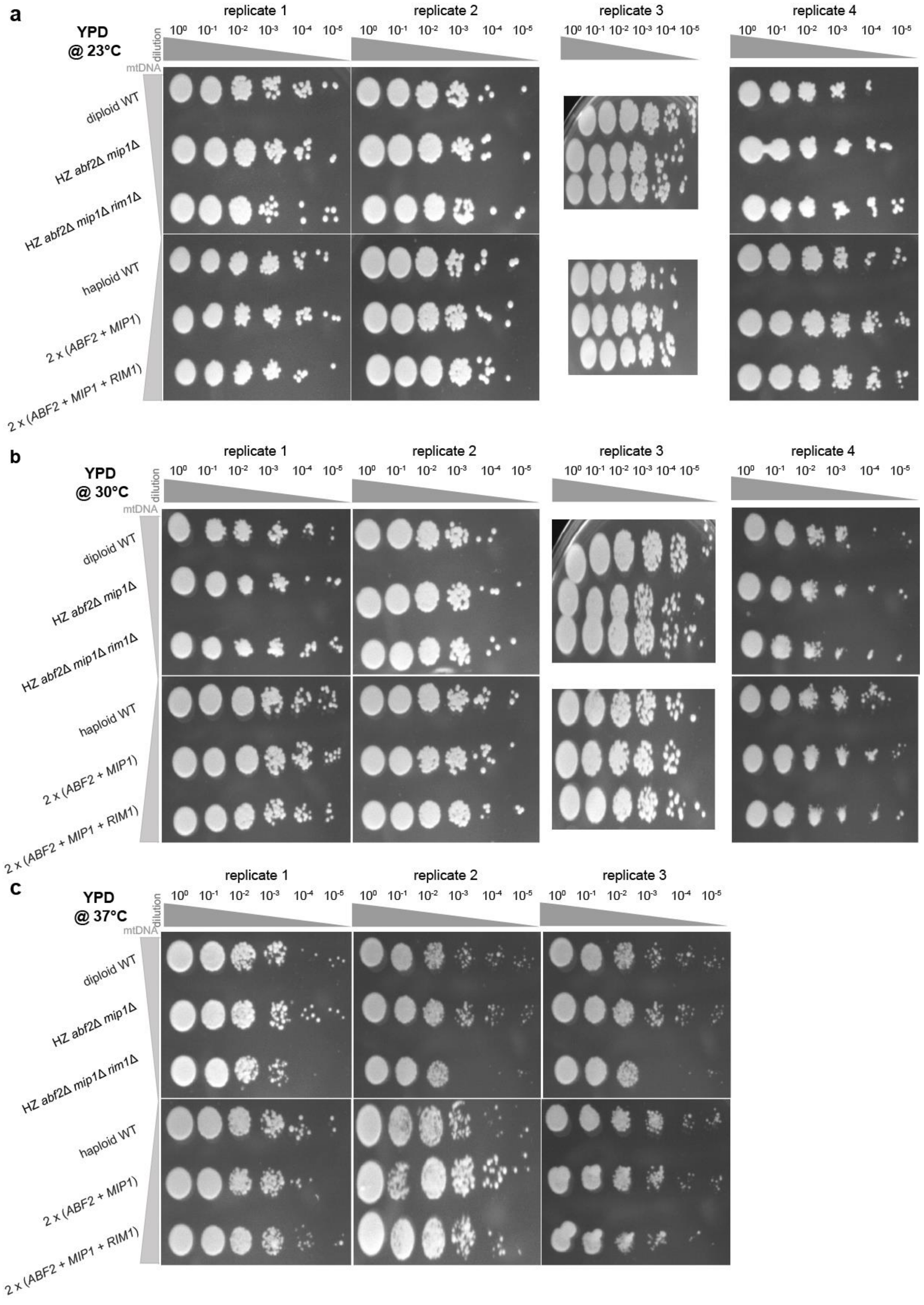
a
YPD
@ 23°C
b
YPD
@ 30°C
c
YPD
@ 37°C
replicate 1
replicate 2
replicate 3
replicate 4
$10^0$ $10^{-1}$ $10^{-2}$ $10^{-3}$ $10^{-4}$ $10^{-5}$
dilution
mtDNA
diploid WT
HZ abf2Δ mip1Δ
HZ abf2Δ mip1Δ rim1Δ
haploid WT
2 x (ABF2 + MIP1)
2 x (ABF2 + MIP1 + RIM1)

**Supplementary Figure S3: mtDNA copy number changes do not lead to major differences in temperature-dependent colony growth on YPD plates.**

Stress spot assays of strains grown on plates with a fermentable carbon source (YPD). Colony growth was tested by plating the strains in a six-step serial dilution, for at least three replicates. Diploids (top) include wild type (WT), a hemizygous deletion of *ABF2* and *MIP1*, as well as a hemizygous deletion of *ABF2, MIP1,* and *RIM1*; haploids (bottom) include WT, a strain with two copies of *ABF2* and *MIP1*, as well as a strain with two copies of *ABF2, MIP1,* and *RIM1*. Plates were incubated for approximately 48 h at different temperatures: **a)** 23 °C; **b)** 30 °C; **c)** 37 °C.

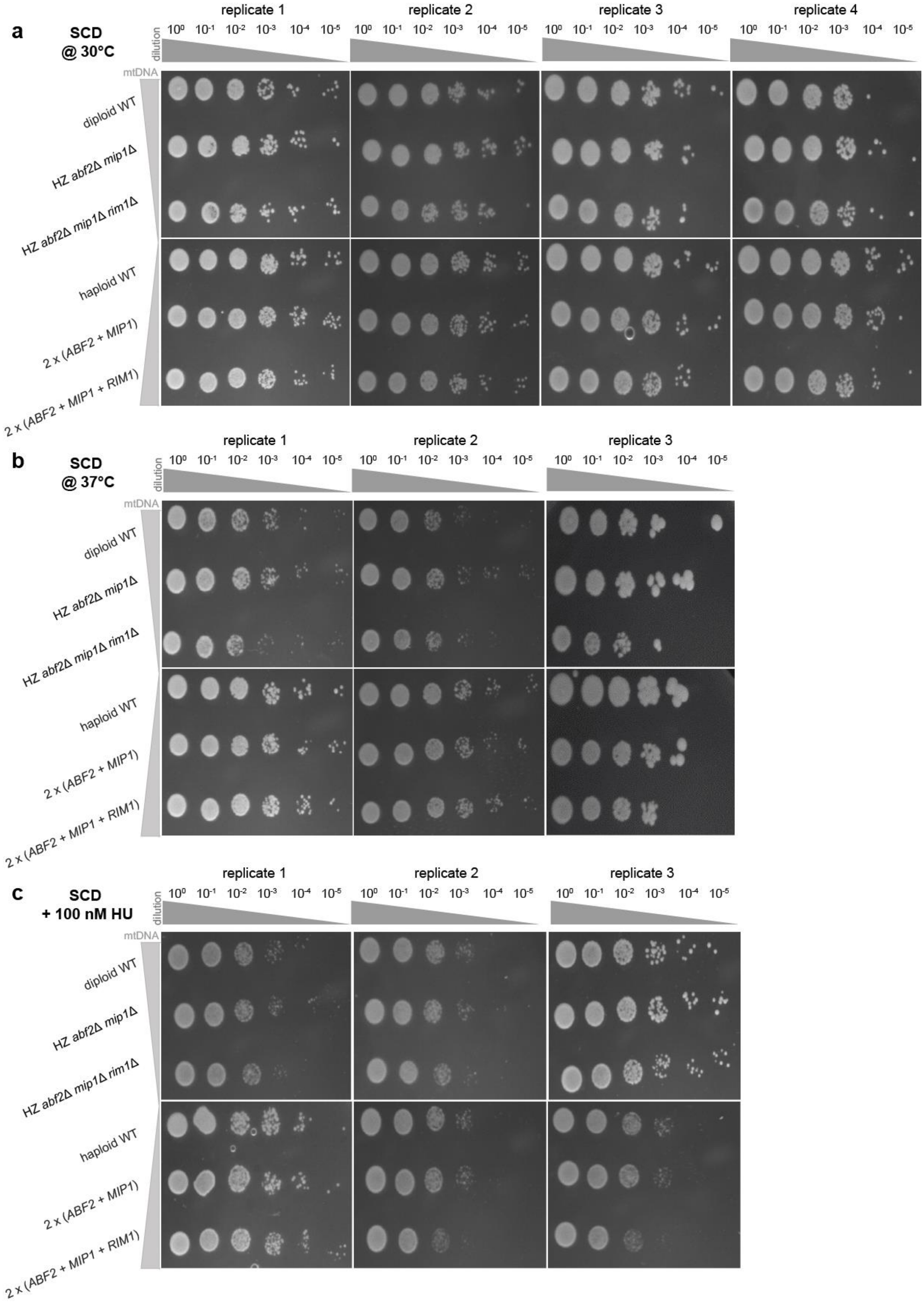

a
SCD @ 30°C
replicate 1
replicate 2
replicate 3
replicate 4
dilution
mtDNA
diploid WT
HZ abf2Δ mip1Δ
HZ abf2Δ mip1Δ rim1Δ
haploid WT
2 x (ABF2 + MIP1)
2 x (ABF2 + MIP1 + RIM1)
b
SCD @ 37°C
c
SCD + 100 nM HU

**Supplementary Figure S4: mtDNA copy number changes do not affect colony growth on SCD plates in increased temperature or upon induction of replicative stress.**

Stress spot assays of strains grown on plates with a fermentable carbon source (SCD). Colony growth was tested by plating the strains in a six-step serial dilution, for at least three replicates. Diploids (top) include wild type (WT), a hemizygous deletion of *ABF2* and *MIP1*, as well as a hemizygous deletion of *ABF2, MIP1,* and *RIM1*; haploids (bottom) include WT, a strain with two copies of *ABF2* and *MIP1*, as well as a strain with two copies of *ABF2, MIP1,* and *RIM1*. Plates were incubated for approximately 48 h at different temperatures (**a)** 30 °C; **b)** 37 °C; **c)** 30 °C) or with supplementation of 100 nM hydroxyurea (c).

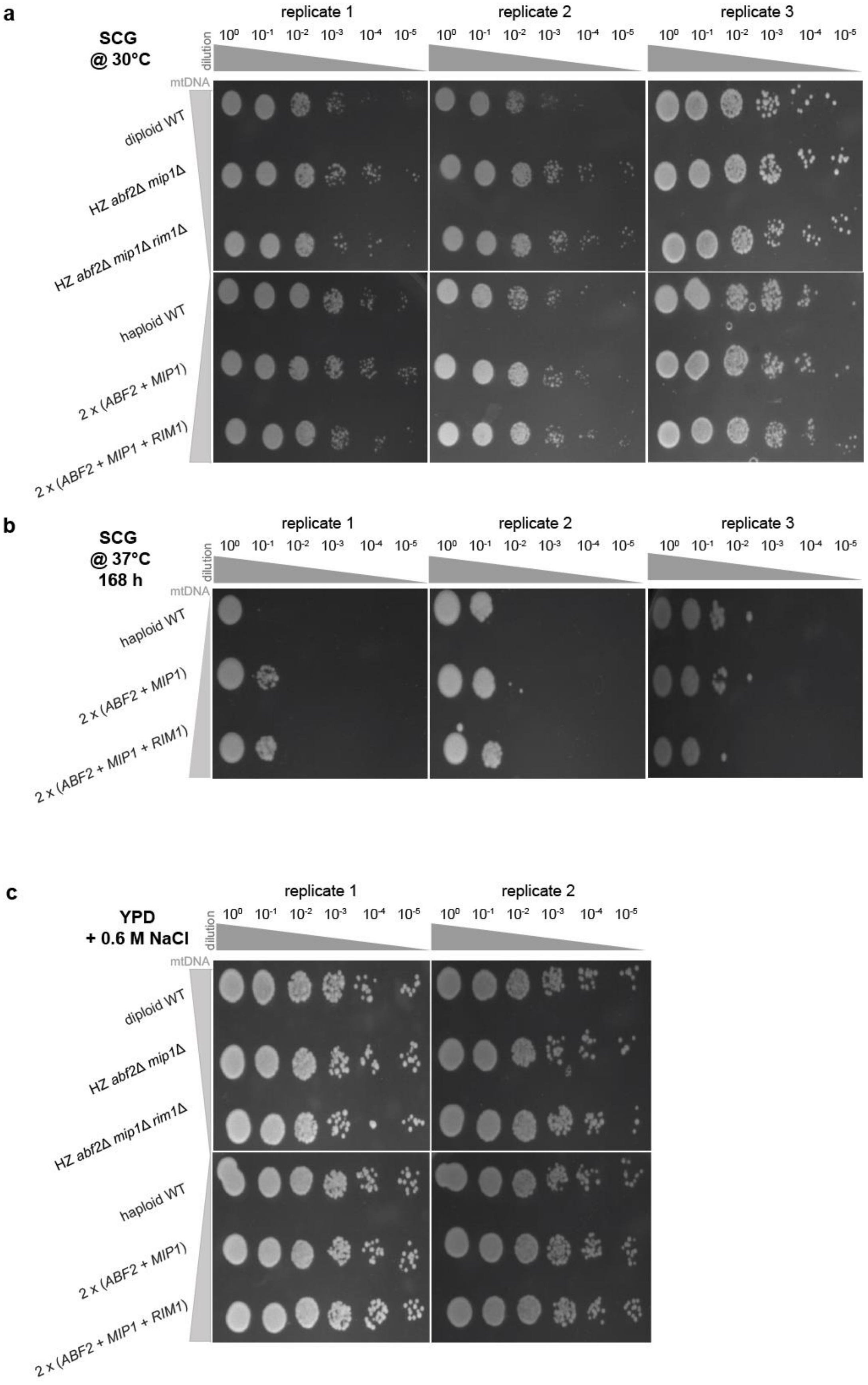
a
SCG
@ 30°C
replicate 1
replicate 2
replicate 3
dilution
10⁰ 10⁻¹ 10⁻² 10⁻³ 10⁻⁴ 10⁻⁵
mtDNA
diploid WT
HZ abf2Δ mip1Δ
HZ abf2Δ mip1Δ rim1Δ
haploid WT
2 x (ABF2 + MIP1)
2 x (ABF2 + MIP1 + RIM1)
b
SCG
@ 37°C
168 h
replicate 1
replicate 2
replicate 3
dilution
10⁰ 10⁻¹ 10⁻² 10⁻³ 10⁻⁴ 10⁻⁵
mtDNA
haploid WT
2 x (ABF2 + MIP1)
2 x (ABF2 + MIP1 + RIM1)
c
YPD
+ 0.6 M NaCl
replicate 1
replicate 2
dilution
10⁰ 10⁻¹ 10⁻² 10⁻³ 10⁻⁴ 10⁻⁵
mtDNA
diploid WT
HZ abf2Δ mip1Δ
HZ abf2Δ mip1Δ rim1Δ
haploid WT
2 x (ABF2 + MIP1)
2 x (ABF2 + MIP1 + RIM1)

**Supplementary Figure S5: mtDNA copy number changes do not affect colony growth at higher temperatures on non-fermentable media or upon osmotic stress in fermentable media.**

Stress spot assays of strains to test colony growth by plating the strains in a six-step serial dilution, for at least two replicates. **a)** Strains were grown on plates with a non-fermentable carbon source (2% glycerol, SCG) and incubated at 30 °C for approximately 48 h. Diploids (top) include wild type (WT), a hemizygous deletion of *ABF2* and *MIP1*, as well as a hemizygous deletion of *ABF2, MIP1,* and *RIM1*; haploids (bottom) include WT, a strain with two copies of *ABF2* and *MIP1*, as well as a strain with two copies of *ABF2, MIP1,* and *RIM1*. **b)** Strains were grown on plates with a non-fermentable carbon source (2% glycerol, SCG) and incubated at 37 °C for 168 h. No colonies grew for any diploid strain tested (not shown); haploids include WT, a strain with two copies of *ABF2* and *MIP1*, as well as a strain with two copies of *ABF2, MIP1,* and *RIM1*. **c)** Strains were grown on plates with a fermentable carbon source (YPD) and additional supplementation of 0.6 M sodium chloride (NaCl) and incubated at 30 °C for approximately 48 h. Diploids (top) include WT, a hemizygous deletion of *ABF2* and *MIP1*, as well as a hemizygous deletion of *ABF2, MIP1,* and *RIM1*; haploids (bottom) include WT, a strain with two copies of *ABF2* and *MIP1*, as well as a strain with two copies of *ABF2, MIP1,* and *RIM1*.

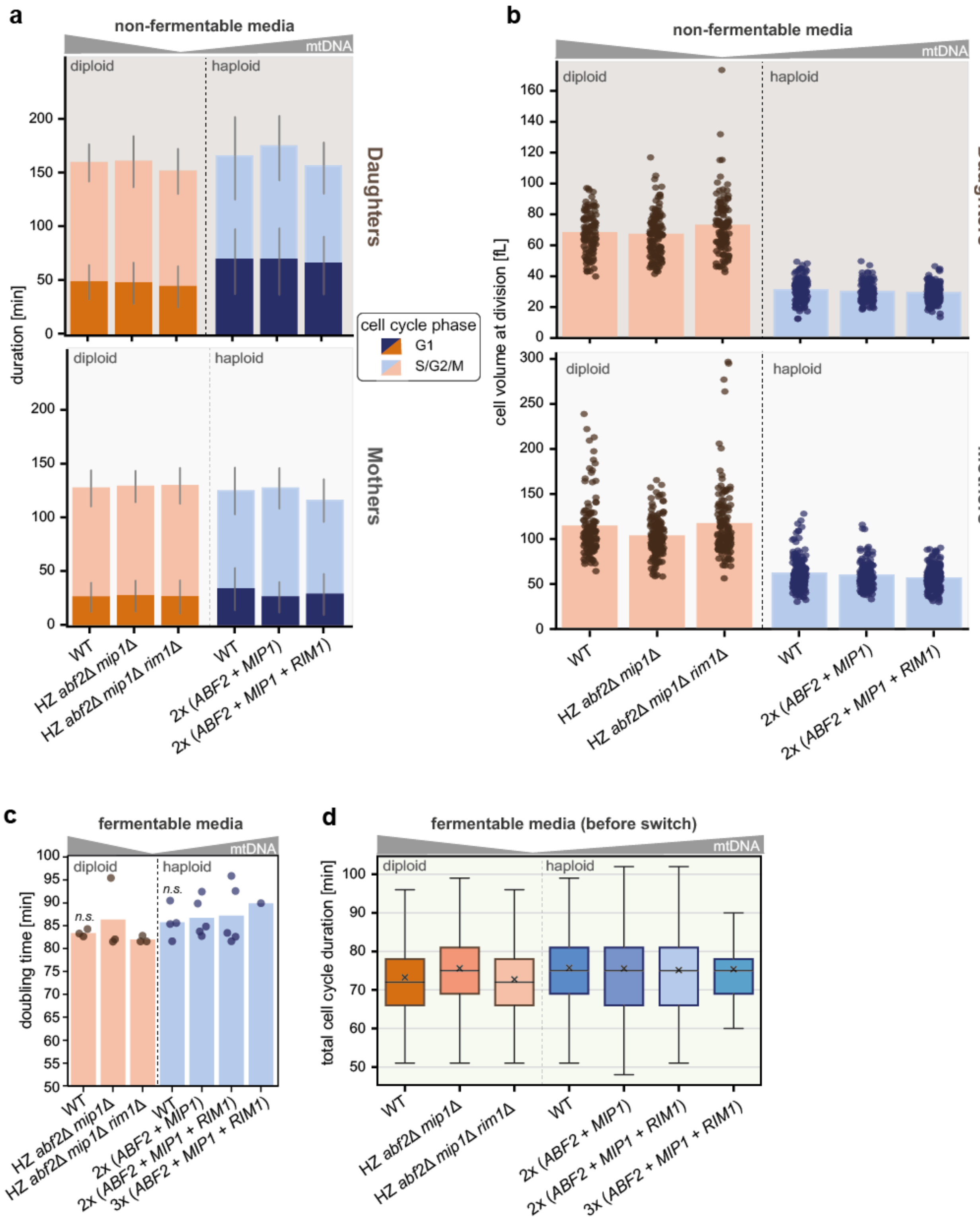

a
non-fermentable media
mtDNA
diploid
haploid
duration [min]
Daughters
Mothers
cell cycle phase
G1
S/G2/M
WT
HZ abf2Δ mip1Δ
HZ abf2Δ mip1Δ rim1Δ
2x (ABF2 + MIP1)
2x (ABF2 + MIP1 + RIM1)
b
cell volume at division [fL]
c
fermentable media
doubling time [min]
n.s.
3x (ABF2 + MIP1 + RIM1)
d
fermentable media (before switch)
total cell cycle duration [min]

**Supplementary Figure S6: Time-lapse microscopy shows that cells are robust to mtDNA copy number changes on a single-cell level.**

**a)** Durations of cell cycle stages for strains grown in non-fermentable media (SCGE) are divided by mother (top; brown background) and daughters (bottom; grey background). Cell cycle stages are categorised into G1 phase (dark) and S/G2/M phase (light). Diploids (orange) include wild type (WT), a hemizygous deletion of *ABF2* and *MIP1*, as well as a hemizygous deletion of *ABF2, MIP1,* and *RIM1*; haploids (blue) include WT, a strain with two copies of *ABF2* and *MIP1*, as well as a strain with two copies of *ABF2, MIP1,* and *RIM1*. Shown are the means of all replicates (bars) and the standard deviation (line). Number of cells measured: $n_{WT\ diploid}$ = 84, $n_{HZ\ abf2\Delta mip1\Delta}$ = 125, $n_{HZ\ abf2\Delta mip1\Delta rim1\Delta}$ = 95, $n_{WT\ haploid}$ = 109, $n_{2x\ (ABF2+MIP1)}$ = 88, $n_{2x\ (ABF2+MIP1+RIM1)}$ = 136.

**b)** Mother (top; brown background) and daughter (bottom; grey background) cell volumes at the time of cell division for strains grown in non-fermentable media (SCGE). Diploids (orange) include WT, a hemizygous deletion of *ABF2* and *MIP1*, as well as a hemizygous deletion of *ABF2, MIP1,* and *RIM1*; haploids (blue) include WT, a strain with two copies of *ABF2* and *MIP1*, as well as a strain with two copies of *ABF2, MIP1,* and *RIM1*. Shown are single replicates (dots) and the mean of all replicates (bars). Numbers of cells measured: $n_{WT\ diploid}$ = 84, $n_{HZ\ abf2\Delta mip1\Delta}$ = 125, $n_{HZ\ abf2\Delta mip1\Delta rim1\Delta}$ = 95, $n_{WT\ haploid}$ = 109, $n_{2x\ (ABF2+MIP1)}$ = 88, $n_{2x\ (ABF2+MIP1+RIM1)}$ = 136.

**c)** Steady state doubling time determined by $OD_{600}$-measurements for various strains grown in fermentable media (SCD). Diploids (orange) include WT, a hemizygous deletion of *ABF2* and *MIP1*, as well as a hemizygous deletion of *ABF2, MIP1,* and *RIM1*; haploids (blue) include WT, a strain with two copies of *ABF2* and *MIP1*, as well as strains with two or three copies of *ABF2, MIP1,* and *RIM1*. Three independent clones were measured and pooled for the strains with two copies of *ABF2* and *MIP1*, as well as strains with two copies of *ABF2, MIP1,* and *RIM1*. Shown are single replicates (dots) and the mean of all replicates (bars). Significance was determined by a two-tailed t-test (* $p<0.05$, ** $p<0.01$, *** $p<0.001$).

**d)** Total cell cycle duration determined in the media-switch time-lapse microscopy experiment, for cells grown in fermentable media (SCD) before the media switch. Diploids (shades of orange) include WT, a hemizygous deletion of *ABF2* and *MIP1*, as well as a hemizygous deletion of *ABF2, MIP1,* and *RIM1*; haploids (shades of blue) include WT, a strain with two copies of *ABF2* and *MIP1*, as well as strains with two or three copies of *ABF2, MIP1,* and *RIM1*. Three independent clones were measured and pooled for the clones with two copies of *ABF2* and *MIP1*, as well as strains with two or three copies of *ABF2, MIP1,* and *RIM1*. Boxplots show the interquartile range (box), total range of data (whisker), median (line), and mean (cross). Number of cells measured: $n_{WTdiploid}$ = 96, $n_{HZ\ abf2\Delta mip1\Delta}$ = 84, $n_{HZ\ abf2\Delta mip1\Delta rim1\Delta}$ = 74, $n_{WT\ haploid}$ = 185, $n_{2x\ (ABF2+MIP1)}$ = 347 (three individual clones pooled), $n_{2x\ (ABF2+MIP1+RIM1)}$ = 281 (three individual clones pooled), $n_{3x\ (ABF2+MIP1+RIM1)}$ = 50.

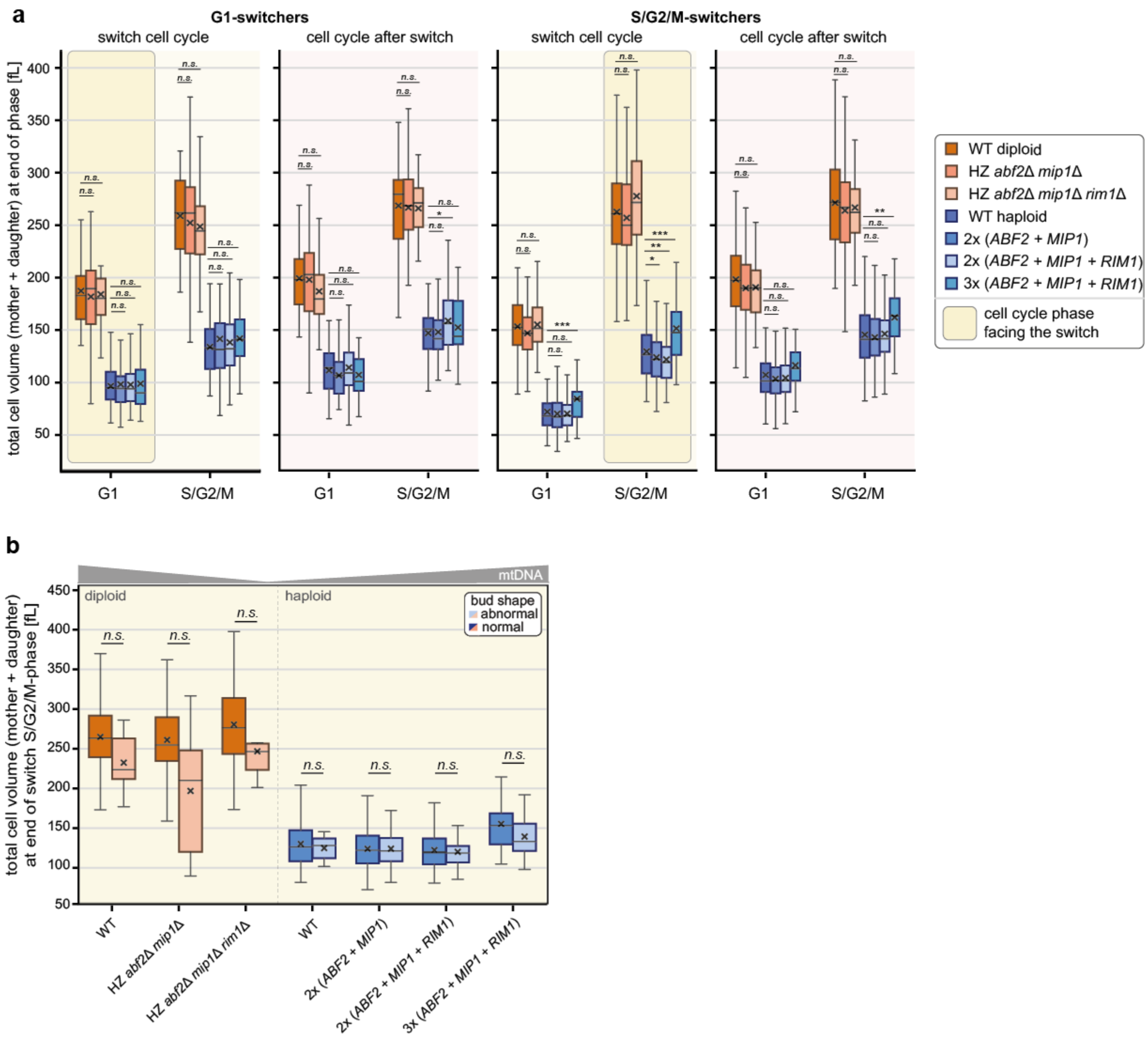


**Supplementary Figure S7: mtDNA copy number changes do not lead to major differences in cell volume of mother and bud during a media switch from normal to low glucose levels.**

**a)** Total cell volumes of mother and daughter at the end of each cell cycle stage are plotted for G1-switchers (2 left panels) and S/G2/M-switchers (2 right panels), highlighting the switch cell cycle phase and the cell cycle after the switch. Boxplots show the interquartile range (box), total range of data (whisker), median (line) and mean (cross). Significance was determined compared to the respective wild type by a two-tailed t-test (* p<0.05, ** p<0.01, *** p<0.001). Number of cells analysed for G1-switchers (left): $n_{\text{WT diploid}}$ = 32 (dark orange), $n_{\text{HZ } abf2\Delta mip1\Delta rim1\Delta}$ = 32 (light orange), $n_{\text{WT haploid}}$ = 83 (dark blue), $n_{\text{2x } (ABF2+MIP1+RIM1)}$ = 122 (light blue; three individual clones pooled), $n_{\text{3x } (ABF2+MIP1+RIM1)}$ = 28 (turquoise), and for S/G2/M-switchers (right): $n_{\text{WT diploid}}$ = 87 (dark orange), $n_{\text{HZ } abf2\Delta mip1\Delta rim1\Delta}$ = 69 (light orange), $n_{\text{WT haploid}}$ = 155 (dark blue), $n_{\text{2x } (ABF2+MIP1+RIM1)}$ = 232 (light blue; three individual clones pooled), $n_{\text{3x } (ABF2+MIP1+RIM1)}$ = 47 (turquoise).

**b)** Total cell volumes of mother and daughter at the end of S/G2/M-phase in the switch cell cycle for normally (dark) and abnormally shaped buds (light), for diploid (orange) and haploid (blue) strains. Boxplots show the interquartile range (box), total range of data (whisker), median (line), and mean (cross). Significance was determined comparing the two types of bud shapes by a two-tailed t-test (* p<0.05, ** p<0.01, *** p<0.001). Number of buds analysed: $n_{\text{WT diploid}}$ = 43, $n_{\text{HZ } abf2\Delta mip1\Delta rim1\Delta}$ = 28, $n_{\text{WT haploid}}$ = 98, $n_{\text{2x } (ABF2+MIP1+RIM1)}$ = 159 (three individual clones pooled), $n_{\text{3x } (ABF2+MIP1+RIM1)}$ = 30.

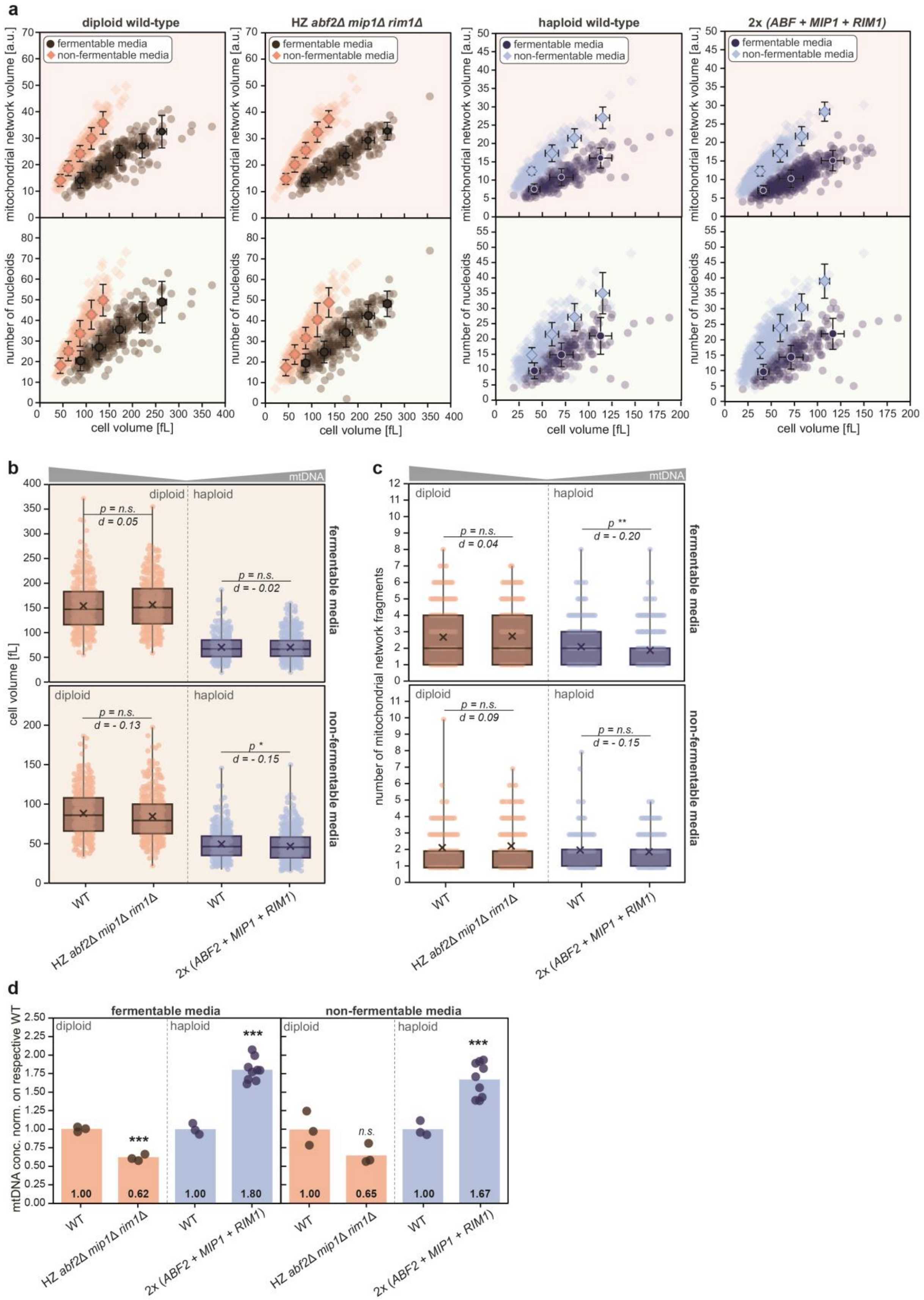

a
diploid wild-type
HZ abf2Δ mip1Δ rim1Δ
haploid wild-type
2x (ABF + MIP1 + RIM1)
fermentable media
non-fermentable media
mitochondrial network volume [a.u.]
number of nucleoids
cell volume [fL]
b
mtDNA
diploid
haploid
p = n.s.
d = 0.05
d = - 0.02
d = - 0.13
p *
d = - 0.15
cell volume [fL]
fermentable media
non-fermentable media
WT
HZ abf2Δ mip1Δ rim1Δ
2x (ABF2 + MIP1 + RIM1)
c
p **
d = 0.04
d = - 0.20
d = 0.09
d = - 0.15
number of mitochondrial network fragments
d
mtDNA conc. norm. on respective WT
***
n.s.
1.00
0.62
1.80
0.65
1.67

**Supplementary Figure S8: Mitochondrial network morphology and nucleoid number depend on cell volume but not mtDNA copy number.**

**a)** Cell volume-dependency of mitochondrial network volume (top; red background) and number of nucleoids (bottom; green background) for various strains grown in fermentable (dark) or non-fermentable (light) media. Shown are single cells (lighter dots), as well as the mean (darker dots) and standard deviations (whiskers) for bins of 50 fL.

**b-c)** Analysis of cell volume (**b**) and the number of mitochondrial network fragments (**c**) from single cells microscopy in fermentable (top) and non-fermentable (bottom) media for diploid (orange) and haploid (blue) strains. Boxplots show the interquartile range (box), total range of data (whisker), median (line), and mean (cross). Number of single cells (dots) measured in three replicates: in fermentable media $n_{\text{WT diploid}}$ = 313, $n_{\text{HZ}\ abf2\Delta mip1\Delta rim1\Delta}$ = 300, $n_{\text{WT haploid}}$ = 347, $n_{2x\ (ABF2+MIP1+RIM1)}$ = 528 (three individual clones pooled); in non-fermentable media: $n_{\text{WT diploid}}$ = 333, $n_{\text{HZ}\ abf2\Delta mip1\Delta rim1\Delta}$ = 331, $n_{\text{WT haploid}}$ = 327, $n_{2x\ (ABF2+MIP1+RIM1)}$ = 581 (three individual clones pooled). Significance was determined compared to the respective wild type using a two-tailed t-test (* p<0.05, ** p<0.01, *** p<0.001), and effect sizes were calculated as Cohen's d.

**d)** mtDNA concentration for microscopy strains grown in fermentable (left) or non-fermentable media (right), normalised on respective wild type, determined by DNA-qPCR. Shown are single replicates (dots) and the mean of all replicates (bars). Significance was determined by a two-tailed t-test (* p<0.05, ** p<0.01, *** p<0.001).

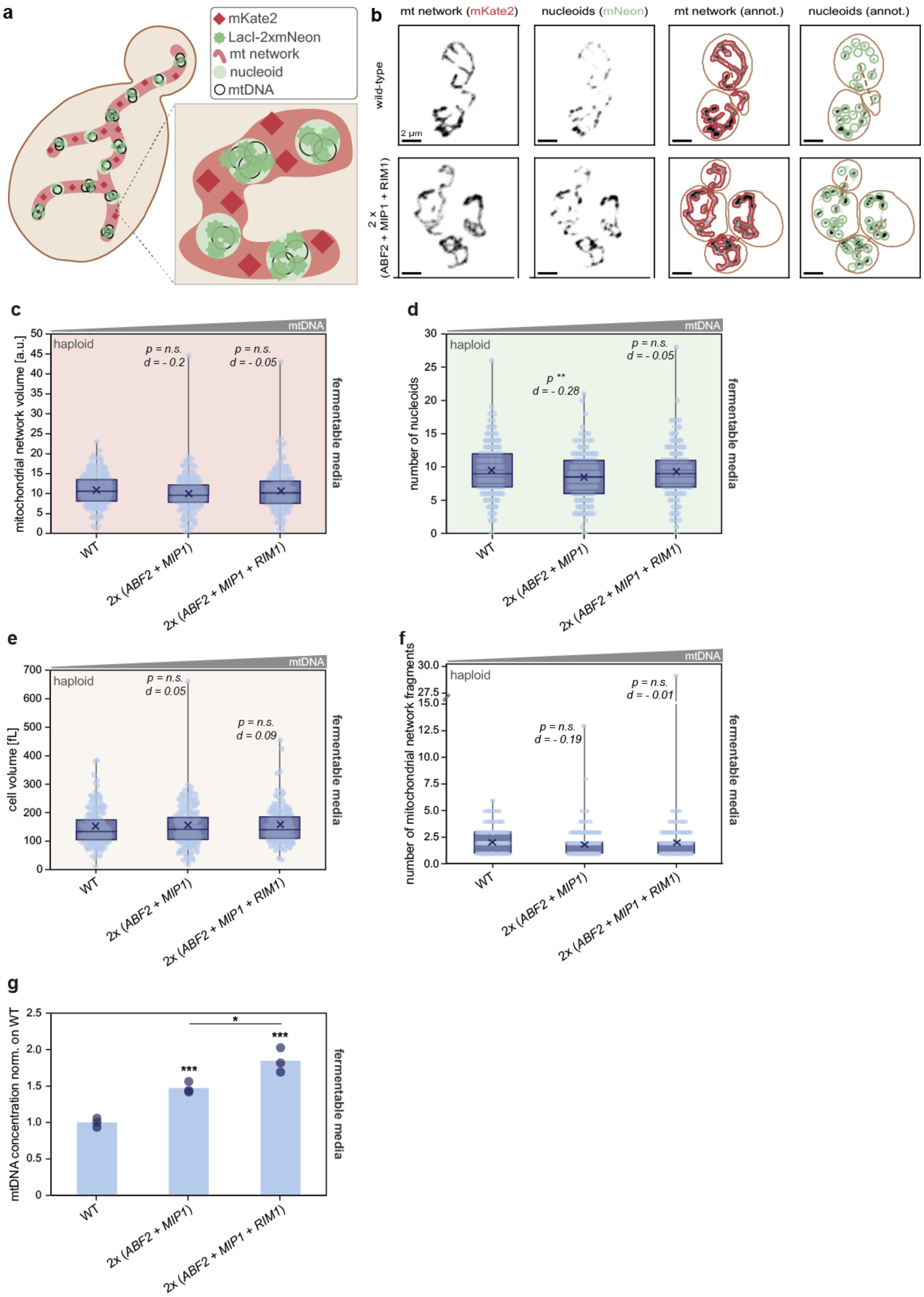
a
mKate2
LacI-2xmNeon
mt network
nucleoid
mtDNA
b
mt network (mKate2)
nucleoids (mNeon)
mt network (annot.)
nucleoids (annot.)
wild-type
2 x (ABF2 + MIP1 + RIM1)
2 µm
c
haploid
mtDNA
p = n.s. d = - 0.2
p = n.s. d = - 0.05
mitochondrial network volume [a.u.]
fermentable media
WT
2x (ABF2 + MIP1)
2x (ABF2 + MIP1 + RIM1)
d
haploid
mtDNA
p ** d = - 0.28
p = n.s. d = - 0.05
number of nucleoids
fermentable media
e
haploid
mtDNA
p = n.s. d = 0.05
p = n.s. d = 0.09
cell volume [fL]
fermentable media
f
haploid
mtDNA
p = n.s. d = - 0.19
p = n.s. d = - 0.01
number of mitochondrial network fragments
fermentable media
g
mtDNA concentration norm. on WT
***
***
*
fermentable media

**Supplementary Figure S9: Alternative visualisation approach confirms that mitochondrial morphology and number of nucleoids are largely independent of mtDNA copy number.**

**a)** Illustration of the LacO-LacI system for mtDNA visualisation. Multiple LacO-repeats have been integrated into the mtDNA[43] and can be bound by mitochondrial-targeted LacI, which is fused to 2xmNeon[5]. The mitochondrial network was visualised with mKate2 targeted to the mitochondrial matrix.

**b)** Representative confocal live-cell images (maximum projections) of the mitochondrial network (mKate2) and nucleoids (mNeon), as well as automatically obtained annotations of mitochondrial network (red), nucleoids (green) and cell segmentations (brown), for cells grown in fermentable media. Strains shown are haploid wild type (top) and a strain with two copies of *ABF2, MIP1,* and *RIM1* (bottom). Scale bars represent 2 µm.

**c-f)** Analysis of **c)** the mitochondrial network volume; **d)** the number of nucleoids; **e)** cell volume; **f)** the number of mitochondrial network fragments from single cells microscopy in fermentable media for haploid strains. Boxplots show the interquartile range (box), total range of data (whisker), median (line) and mean (cross). Number of single cells (dots) measured in three replicates: $n_{\text{WT haploid}}$ = 176, $n_{2x\ (ABF2+MIP1+RIM1)}$ = 178, $n_{2x\ (ABF2+MIP1+RIM1)}$ = 163. Significance was determined compared to wild type using a two-tailed t-test (* p<0.05, ** p<0.01, *** p<0.001), and effect sizes were calculated as Cohen's d.

**g)** mtDNA concentration for microscopy strains grown in fermentable media, normalised on haploid wild type, determined by DNA-qPCR. Shown are single replicates (blue dots) and the mean of all replicates (bars). Significance was determined by a two-tailed t-test (* p<0.05, ** p<0.01, *** p<0.001).

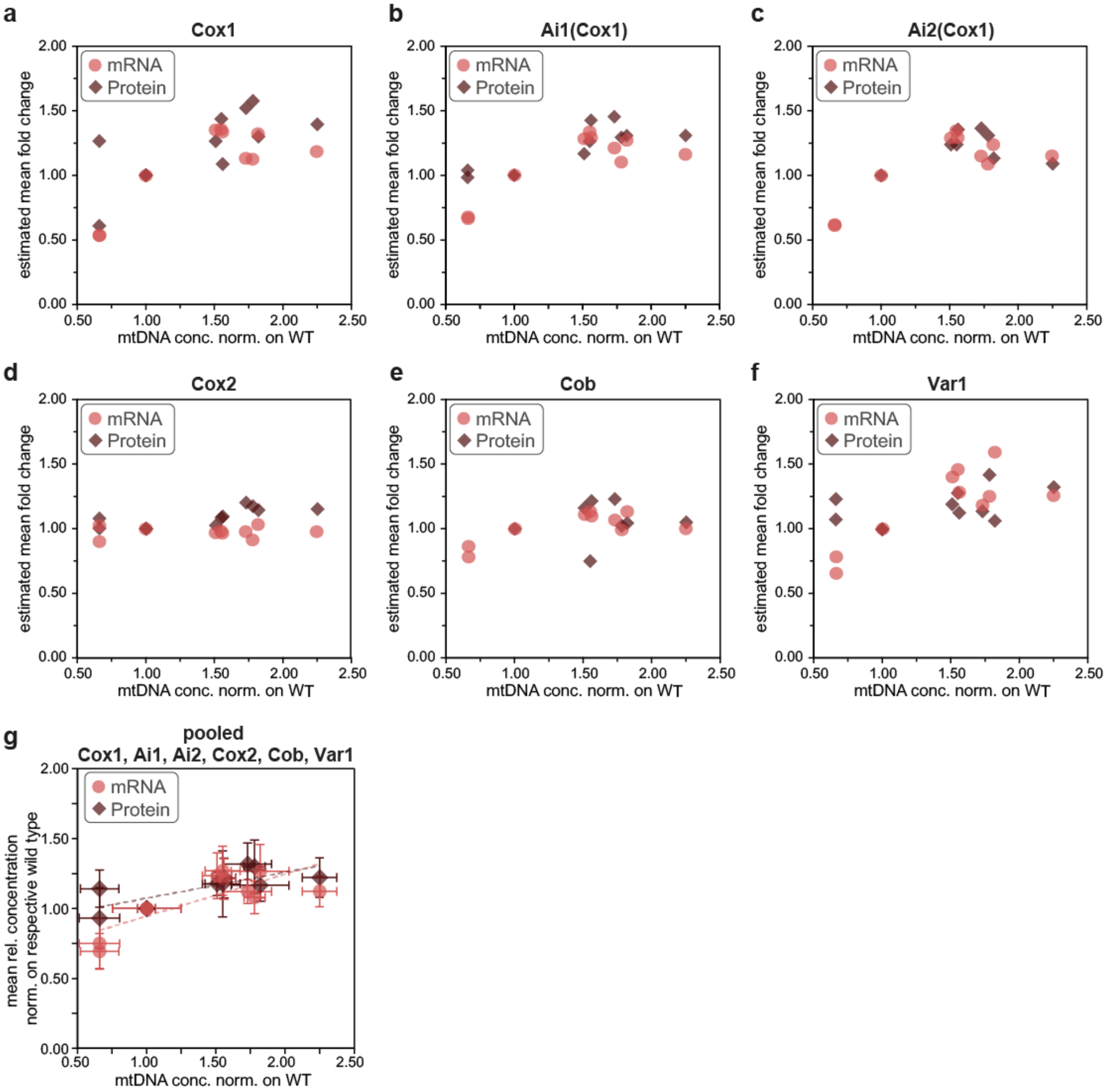


**Supplementary Figure S10: mRNA levels corresponding to the detected proteins depend more strongly on mtDNA copy number than the relative protein concentrations.**

Estimated mean fold change as a function of the mtDNA concentration, both normalised on the respective wild type on mRNA-level (light red dot) and protein-level (dark red diamond). Means represent at least three replicates for mRNA and two replicates for proteins. mtDNA-dependency shown for: Cox1 (**a**), Ai1(Cox1) (**b**), Ai2(Cox1) (**c**), Cox2 (**d**), Cob (**e**), Var1 (**f**), and the mean behaviour for all 6 detected proteins (**g**).

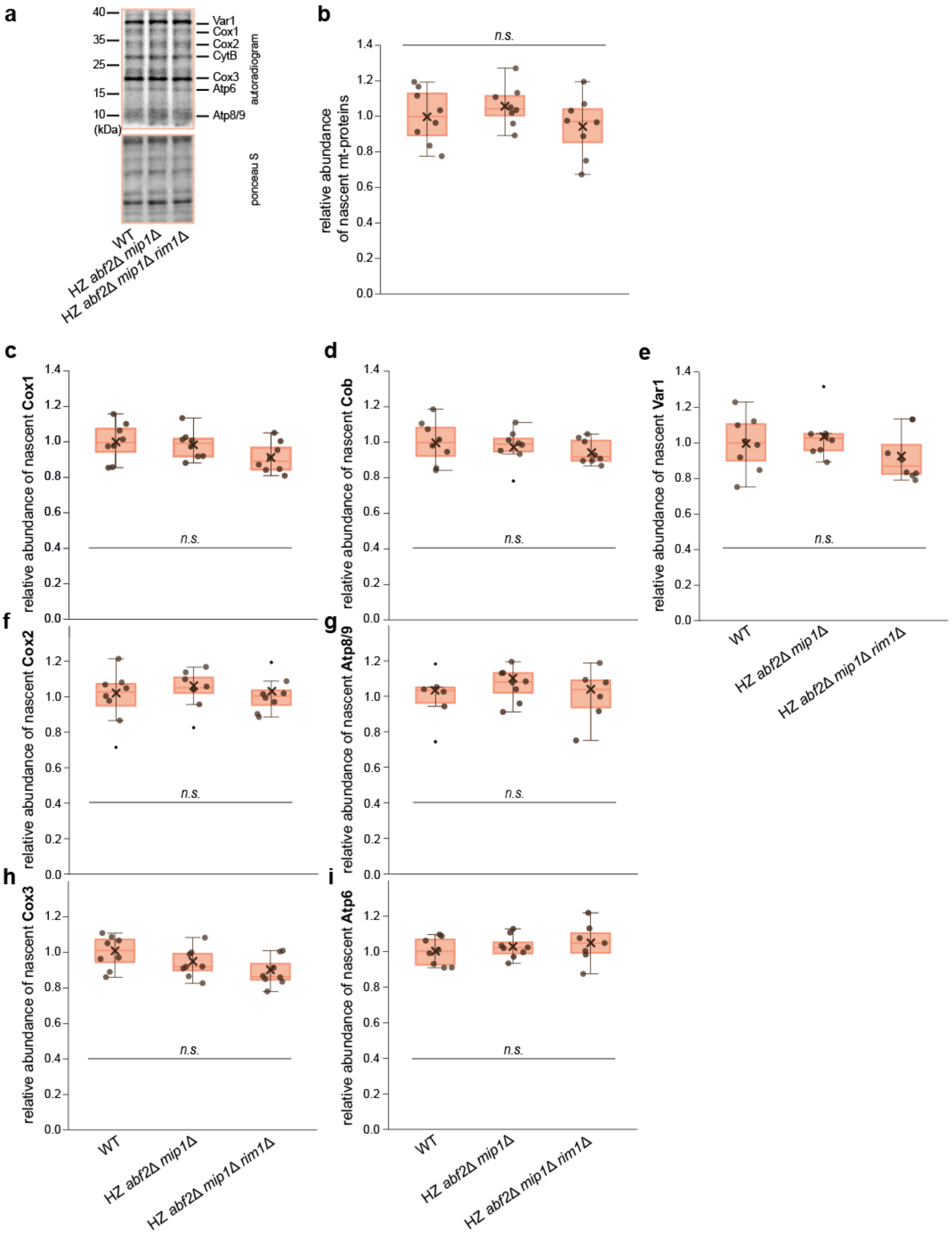


**Supplementary Figure S11: Decreasing mtDNA copy numbers in diploid strains do not affect the abundance of individual nascent mtDNA-encoded proteins.**

**a)** Example of autoradiogram (top) and corresponding Ponceau S staining (bottom).

**b-i)** Relative abundance of nascent mtDNA-encoded proteins determined by radiolabelling experiment. Boxplots show the interquartile range (box), total range of data (whisker), median (line), and mean (cross). Single replicates are shown as dots. Diploids (orange) include wild type (WT), a hemizygous deletion of *ABF2* and *MIP1*, as well as a hemizygous deletion of *ABF2, MIP1,* and *RIM1. S*tatistical significance was evaluated using one-way ANOVA with Tukey's post hoc test, or, when assumptions were not met, a Kruskal–Wallis test followed by pairwise Wilcoxon tests with Benjamini–Hochberg correction. Relative abundance of total nascent mtDNA-encoded proteins (**b**), or individual proteins: Cox1 (**c**), Cob (**d**), Var1 (**e**), Cox2 (**f**), Atp8/9 (**g**), Cox3 (**h**), Atp6 (**i**).

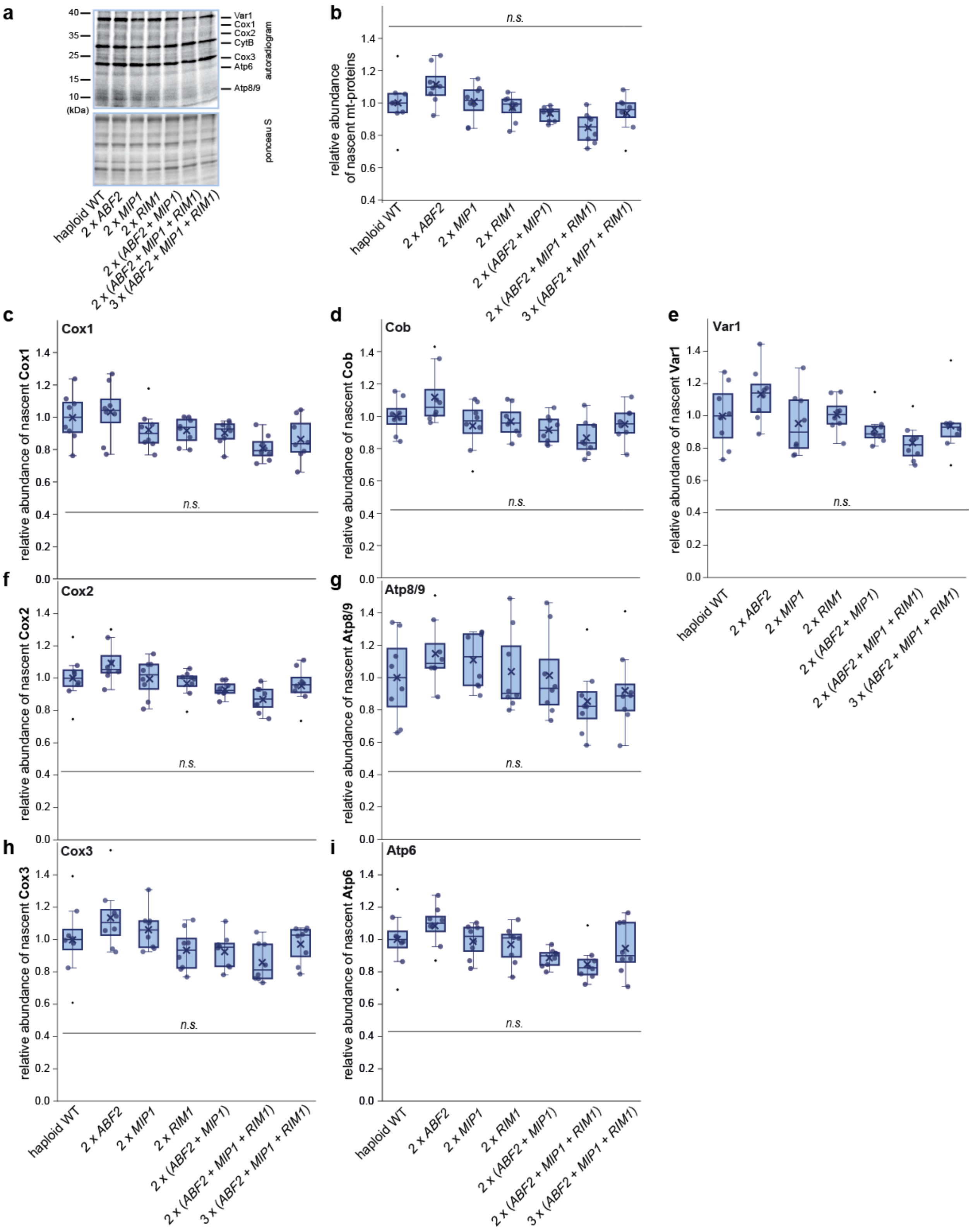
a
Var1
Cox1
Cox2
CytB
Cox3
Atp6
Atp8/9
autoradiogram
ponceau S
(kDa)
haploid WT
2 x ABF2
2 x MIP1
2 x RIM1
2 x (ABF2 + MIP1)
2 x (ABF2 + MIP1 + RIM1)
3 x (ABF2 + MIP1 + RIM1)
b
relative abundance of nascent mt-proteins
n.s.
c
Cox1
relative abundance of nascent Cox1
d
Cob
relative abundance of nascent Cob
e
Var1
relative abundance of nascent Var1
f
Cox2
relative abundance of nascent Cox2
g
Atp8/9
relative abundance of nascent Atp8/9
h
Cox3
relative abundance of nascent Cox3
i
Atp6
relative abundance of nascent Atp6

**Supplementary Figure S12: Increasing mtDNA copy numbers in haploid strains do not affect the abundance of individual nascent mtDNA-encoded proteins.**

**a)** Example of autoradiogram (top) and corresponding Ponceau S staining (bottom).

**b-i)** Relative abundance of nascent mtDNA-encoded proteins determined by radiolabelling experiment. Boxplots show the interquartile range (box), total range of data (whisker), median (line), and mean (cross). Single replicates are shown as dots. Haploids (blue) include wild type (WT), strains with two copies of *ABF2*, or *MIP1*, or *RIM1*, as well as a strain with two copies of *ABF2* and *MIP1*, and strains with two or three copies of *ABF2, MIP1,* and *RIM1*. *S*tatistical significance was evaluated using one-way ANOVA with Tukey's post hoc test, or, when assumptions were not met, a Kruskal–Wallis test followed by pairwise Wilcoxon tests with Benjamini–Hochberg correction. Relative abundance of total nascent mtDNA-encoded proteins (**b**), or individual proteins: Cox1 (**c**), Cob (**d**), Var1 (**e**), Cox2 (**f**), Atp8/9 (**g**), Cox3 (**h**), Atp6 (**i**).

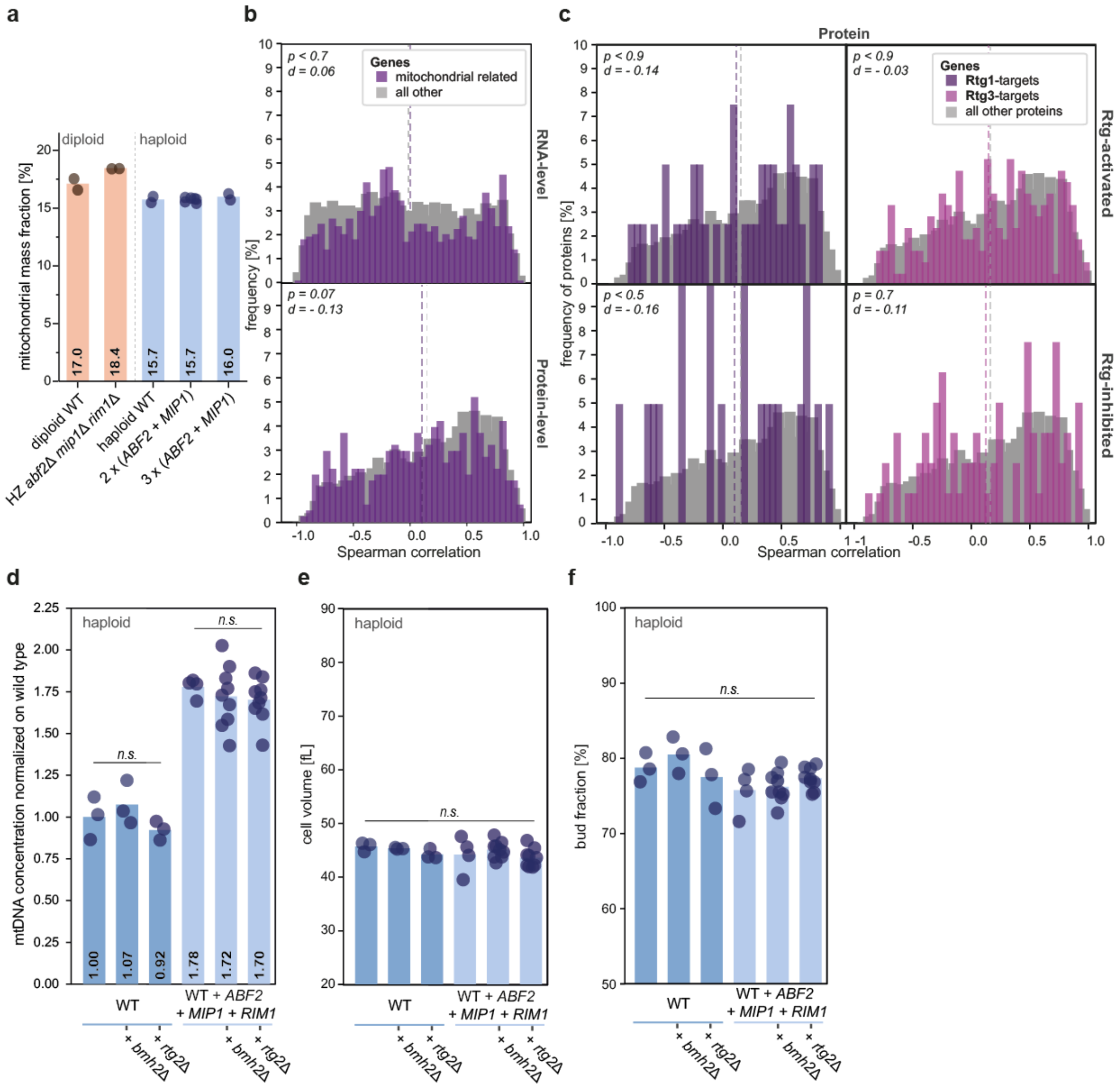


**Supplementary Figure S13: Interplay of retrograde signalling and mtDNA copy number.**

**a)** Mitochondrial mass fraction estimated by mass spectrometry analysis shows no difference with altered mtDNA copy numbers. Shown are single replicates (dots) and the mean of all replicates (bars). Three independent clones were measured and pooled for the strain with two copies of *ABF2, MIP1,* and *RIM1*. Only the proteins that are exclusively located in mitochondria were included in the estimation of mitochondrial mass fraction.

**b)** Histogram of Spearman correlation coefficients for mitochondria-related genes (purple) and all other genes (grey). Separated into results on RNA- (top) and protein-level (bottom). The mean of each category is shown as a dashed line. The significance between distributions was determined by a two-sided Fisher's Exact test (* p<0.05, ** p<0.01, *** p<0.001), and effect sizes were calculated as Cohen's d.

**c)** Histogram of Spearman correlation coefficients for Rtg1- or Rtg3-dependent proteins (purple and pink) and all other proteins (grey). Separated by proteins that are activated (top) or inhibited (bottom). The mean of each category is shown as a dashed line. The significance between distributions was determined by a two-sided Fisher's Exact test (* p<0.05, ** p<0.01, *** p<0.001), and effect sizes were calculated as Cohen's d.

**d-f)** Measurements of **d)** mtDNA concentration normalised on haploid WT by DNA-qPCR; **e)** Mean cell volume determined by Coulter counter; **f)** Percentage of bud fraction determined by visual inspection under the microscope for various strains grown in non-fermentable media. Haploid strains include WT and a strain with two copies of *ABF2, MIP1,* and *RIM1*, both of which were either unmodified or harbouring additional deletions of *BMH2* or *RTG2*. Shown are single replicates (dots) and the mean of all replicates (bars). Significance was determined by a two-tailed t-test (* p<0.05, ** p<0.01, *** p<0.001).

**Supplementary Table 1: Strains used in this study.**

Note that strains with additional copies of genes include 1000 bp upstream and at least 200 bp downstream of the coding sequence.

| Name | Genotype | Description | Origin | Figure |
|---|---|---|---|---|
| AFY003-1 | *Mat a/α; ADE2/ADE2 URA3/ura3, LEU2/leu2 mip1::CglaTRP1/MIP1 abf2::clonNAT/ABF2 rim1::HIS3/RIM1* | Triple hemizygous deletion of *mip1, abf2, and rim1* | This study | 1 b, d, e, g; 2 a-c, e, g, i, j; 4 a-d, f, g; 5 a-c, e, f; S 1 a, b, d, h; S 2 a, b; S 3-7; S 10; S.11; S 13 a-c |
| AFY004-3 | *Mat a/α; ADE2/ADE2 URA3/ura3, LEU2/leu2 mip1::CglaTRP1/MIP1 abf2::clonNAT/ABF2 mhr1::HIS3/MHR1* | Triple hemizygous deletion of *mip1, abf2, and mhr1* | This study | 1 b; S1 a |
| AFY005-1 | *Mat a/α; ADE2/ADE2 URA3/ura3, LEU2/leu2 mip1::CglaTRP1/MIP1 abf2::clonNAT/ABF2 hmi1::HIS3/HMI1* | Triple hemizygous deletion of *mip1, abf2, and hmi1* | This study | 1 b; S1 a |
| AFY006-3 | *Mat a/α; ADE2/ADE2 URA3/ura3, LEU2/leu2 mip1::CglaTRP1/MIP1 abf2::clonNAT/ABF2 mgm101::HIS3/MGM101* | Triple hemizygous deletion of *mip1, abf2, and mgm101* | This study | 1 b; S1 a |
| AFY007-7 | *Mat a/α; ADE2/ADE2 URA3/ura3, LEU2/leu2 mip1::CglaTRP1/MIP1 abf2::clonNAT/ABF2 pif1::HIS3/PIF1* | Triple hemizygous deletion of *mip1, abf2, and pif1* | This study | 1 b; S1 a |
| AFY008-1 | *Mat a/α; ADE2/ADE2 URA3/ura3, LEU2/leu2 mip1::CglaTRP1/MIP1 abf2::clonNAT/ABF2 rpo41::HIS3/RPO41* | Triple hemizygous deletion of *mip1, abf2, and rpo41* | This study | 1 b; S1 a |
| AFY009-2 | *Mat a/α; ADE2/ADE2 URA3/ura3, LEU2/leu2 mip1::CglaTRP1/MIP1 abf2::clonNAT/ABF2 mtf1::HIS3/MTF1* | Triple hemizygous deletion of *mip1, abf2, and mtf1* | This study | 1 b; S1 a |
| AFY011-1 | *Mat a/α; ADE2/ADE2 URA3/ura3, LEU2/leu2 abf2::clonNAT/ABF2 rim1::HIS3/RIM1* | Double hemizygous deletion of *abf2, and rim1* | This study | 1 b; S1 a |
| AFY012-1 | *Mat a/α; ADE2/ADE2 URA3/ura3, LEU2/leu2* | Double hemizygous | This study | 1 b; S1 a |

| | | | | |
|---|---|---|---|---|
| | *abf2::clonNAT/ABF2 mhr1::HIS3/MHR1* | deletion of *abf2*, and *mhr1* | | |
| AFY013-1 | *Mat a/α; ADE2/ADE2 URA3/ura3, LEU2/leu2 abf2::clonNAT/ABF2 hmi1::HIS3/HMI1* | Double hemizygous deletion of *abf2*, and *hmi1* | This study | 1 b; S1 a |
| AFY014-1 | *Mat a/α; ADE2/ADE2 URA3/ura3, LEU2/leu2 abf2::clonNAT/ABF2 mgm101::HIS3/MGM101* | Double hemizygous deletion of *abf2*, and *mgm101* | This study | 1 b; S1 a |
| AFY015-1 | *Mat a/α; ADE2/ADE2 URA3/ura3, LEU2/leu2 abf2::clonNAT/ABF2 pif1::HIS3/PIF1* | Double hemizygous deletion of *abf2*, and *pif1* | This study | 1 b; S1 a |
| AFY016-1 | *Mat a/α; ADE2/ADE2 URA3/ura3, LEU2/leu2 abf2::clonNAT/ABF2 rpo41::HIS3/RPO41* | Double hemizygous deletion of *abf2*, and *rpo41* | This study | 1 b; S1 a |
| AFY017-1 | *Mat a/α; ADE2/ADE2 URA3/ura3, LEU2/leu2 abf2::clonNAT/ABF2 mtf1::HIS3/MTF1* | Double hemizygous deletion of *abf2*, and *mtf1* | This study | 1 b; S1 a |
| AFY018-1 | *Mat a/α; ADE2/ADE2 URA3/ura3, LEU2/leu2 mip1::CglaTRP1/MIP1 rim1::HIS3/RIM1* | Double hemizygous deletion of *mip1*, and *rim1* | This study | 1 b; S1 a |
| AFY019-1 | *Mat a/α; ADE2/ADE2 URA3/ura3, LEU2/leu2 mip1::CglaTRP1/MIP1 mhr1::HIS3/MHR1* | Double hemizygous deletion of *mip1*, and *mhr1* | This study | 1 b; S1 a |
| AFY020-1 | *Mat a/α; ADE2/ADE2 URA3/ura3, LEU2/leu2 mip1::CglaTRP1/MIP1 hmi1::HIS3/HMI1* | Double hemizygous deletion of *mip1*, and *hmi1* | This study | 1 b; S1 a |
| AFY021-1 | *Mat a/α; ADE2/ADE2 URA3/ura3, LEU2/leu2 mip1::CglaTRP1/MIP1 mgm101::HIS3/MGM101* | Double hemizygous deletion of *mip1*, and *mgm101* | This study | 1 b; S1 a |
| AFY022-1 | *Mat a/α; ADE2/ADE2 URA3/ura3, LEU2/leu2 mip1::CglaTRP1/MIP1 pif1::HIS3/PIF1* | Double hemizygous deletion of *mip1*, and *pif1* | This study | 1 b; S1 a |
| AFY023-1 | *Mat a/α; ADE2/ADE2 URA3/ura3, LEU2/leu2 mip1::CglaTRP1/MIP1 rpo41::HIS3/RPO41* | Double hemizygous deletion of *mip1*, and *rpo41* | This study | 1 b; S1 a |
| AFY024-1 | *Mat a/α; ADE2/ADE2 URA3/ura3, LEU2/leu2 mip1::CglaTRP1/MIP1 mtf1::HIS3/MTF1* | Double hemizygous deletion of *mip1*, and *mtf1* | This study | 1 b; S1 a |

| | | | | |
|---|---|---|---|---|
| AFY025-1 | *Mat a/α; ADE2/ADE2 URA3/ura3, LEU2/leu2 mip1::CglaTRP1/MIP1 abf2::clonNAT/ABF2 rim1::HIS3/RIM1 hmi1::KanMX6/HMI1* | Quadruple hemizygous deletion of *abf2, mip1, rim1, and hmi1* | This study | 1 b, d;<br>S1 a, d |
| AFY033-1 | *Mat α; ADE2 trp1::MIP1-TRP1 his3::HMI1-HIS3* | Haploid WT with an additional copy of *MIP1,* and *HMI1* | This study | 1 c;<br>S1 c |
| AFY034-2 | *Mat α; ADE2 ura3::ABF2-URA3 trp1::MIP1-TRP1 his3::HMI1-HIS3* | Haploid WT an additional copy of *ABF2*, *MIP1,* and *HMI1* | This study | 1 c;<br>S1 c |
| AFY037-1 | *Mat α; ADE2 leu2::RIM1-LEU2* | Haploid WT with an additional copy of *RIM1* | This study | 1 c;<br>S1 c, g |
| AFY037-2 | *Mat α; ADE2 leu2::2xRIM1-LEU2* | Haploid WT with two additional copies of *RIM1* | This study | S1 g |
| AFY037-3 | *Mat α; ADE2 leu2::RIM1-LEU2* | Haploid WT with an additional copy of *RIM1* | This study | 1 c;<br>S1 c, g |
| AFY053-2 | *Mat α; ADE2 ura3::ABF2-URA3 his3::HMI1-HIS3* | Haploid WT with an additional copy of ABF2, and *HMI1* | This study | 1 c;<br>S1 c |
| AFY054-4 | *Mat α; ADE2 his3::HMI1-HIS3* | Haploid WT with an additional copy of *HMI1* | This study | 1 c;<br>S1 c, g |
| AFY054-7 | *Mat α; ADE2 his3::2xHMI1-HIS3* | Haploid WT with two additional copies of *HMI1* | This study | S1 g |
| AFY054-9 | *Mat α; ADE2 his3::3xHMI1-HIS3* | Haploid WT with three additional copies of *HMI1* | This study | S1 g |
| AFY055-5 | *Mat α; ADE2 ura3::ABF2-URA3 trp1::MIP1-TRP1 leu2::RIM1-LEU2 his3::HMI1-HIS3* | Haploid WT with an additional copy of *ABF2, MIP1, RIM1* and *HMI1* | This study | 1 d;<br>S1 d |
| AFY056-2 | *Mat α; ADE2 ura3::ABF2-URA3 leu2::RIM1-LEU2* | Haploid WT with an additional copy of *ABF2,* and *RIM1* | This study | 1 c;<br>S1 c |
| AFY057-1 | *Mat α; ADE2 ura3::ABF2-URA3 trp1::MIP1-TRP1 leu2::RIM1-LEU2* | Haploid WT with an additional copy of *ABF2, MIP1,* and *RIM1* | This study | 1 c, d, e, g;<br>2 a-e, g, i, j;<br>4 a-d, f, g;<br>5 a-c, e, f;<br>S1 c, d, h; |

| | | | | |
|---|---|---|---|---|
| | | | | S2 a, b; S3-7; S 10; S 12; S13 a-c |
| AFY058-2 | *Mat α; ADE2 leu2::3xRIM1-LEU2* | Haploid WT with three additional copies of *RIM1* | This study | S1 g |
| AFY059-2 | *Mat α; ADE2 leu2::4xRIM1-LEU2* | Haploid WT with four additional copies of *RIM1* | This study | S1 g |
| AFY060 | *Mat α; ADE2 trp1::MIP1-TRP1 leu2::RIM1-LEU2* | Haploid WT with an additional copy of *MIP1,* and *RIM1* | This study | 1 c; S1 c |
| AFY067-1 | *Mat α; ADE2 his3::LexA-ER-AD-TF-HIS3 ura3::LexApr-ABF2-CYC1term-URA3 abf2::hphMX* | Abf2-inducible strain with additional deletion of endogenous *abf2* | This study | S1 e-f |
| AFY070-5 | *Mat a; ADE2 TRP1 mt-LacO CUP1pr-SU9-2xmNEON-LACI-PGK1pr-SU9-mKATE2:KanMX4 ura3::ABF2-URA3* | Haploid microscopy strain with an additional copy of *ABF2* | This study | S9 |
| AFY072-1 | *Mat α; ADE2 TRP1 URA3 LEU2 HIS3* | Haploid WT corrected auxotrophic markers of *TRP1*, *URA3*, *LEU2,* and *HIS3* | This study | S2 d |
| AFY074-1 | *Mat α; ADE2 ura3::2xABF2-URA3 trp1::MIP1-TRP1 his3::MIP1-HIS3 leu2::2xRIM1-LEU2* | Haploid WT with two additional copies of *ABF2, MIP1,* and *RIM1* | This study | 1 e, g; 2 a-c, e, g, i, j; 4 a-d, f, g; 5 a-c, e, f; S1 d, h; S2 a, b, d; S6 c-d; S7; S 10; S 12; 13 a-c |
| AFY082-2 | *Mat α; ADE2 ura3::ABF2-URA3 trp1::MIP1-TRP1 his3::MGM101-HIS3* | Haploid WT with an additional copy of *ABF2, MIP1,* and *MGM101* | This study | 1 c; S1 c |
| AFY083-1 | *Mat α; ADE2 ura3::ABF2-URA3 trp1::MIP1-TRP1* | Haploid WT with an additional copy of *ABF2,* | This study | 1 d; S1 d |

| | | | | |
|---|---|---|---|---|
| | *leu2::RIM1-LEU2*<br>*his3::MGM101-HIS3* | *MIP1, RIM1,*<br>and *MGM101* | | |
| AFY086-1 | *Mat α; ADE2 TRP1 URA3* | Haploid WT corrected auxotrophic markers of *TRP1* and *URA3* | This study | S2 c |
| AFY092-1 | *Mat α; ADE2 TRP1 URA3 LEU2* | Haploid WT corrected auxotrophic markers of *TRP1*, *URA3,* and *LEU2* | This study | 2 d |
| AFY098-1 | *Mat α; ADE2*<br>*ura3::ABF2-URA3*<br>*trp1::MIP1-TRP1* | Haploid WT with an additional copy of *ABF2,* and *MIP1* | This study | 1 a, c, e;<br>2 e;<br>4 a-d, f, g;<br>5 a-c, e, f;<br>S1 c; S2 a-c;<br>S3-7; S10;<br>S12; S13 a-b |
| AFY098-2 | *Mat α; ADE2*<br>*ura3::ABF2-URA3*<br>*trp1::MIP1-TRP1* | Haploid WT with an additional copy of *ABF2,* and *MIP1* | This study | 1 a, c, e;<br>2 e;<br>4 a-d, f, g;<br>5 a-c, e, f;<br>S1 c; S2 a-c;<br>S3-7; S10;<br>S12; S13 a-b |
| AFY099-1 | *Mat a; ADE2*<br>*TRP1*<br>*mt-LacO*<br>*CUP1pr-SU9-2xmNEON-LACI-PGK1pr-SU9-mKATE2:KanMX4*<br>*ura3::ABF2-URA3*<br>*his3::MIP1-HIS3* | Haploid microscopy strain with an additional copy of *ABF2,* and *MIP1* | This study | S9 |
| AFY100-1 | *Mat a; ADE2*<br>*TRP1*<br>*mt-LacO*<br>*CUP1pr-SU9-2xmNEON-LACI-PGK1pr-SU9-mKATE2:KanMX4*<br>*ura3::ABF2-URA3*<br>*his3::MIP1-HIS3*<br>*leu2::RIM1-LEU2* | Haploid microscopy strain with an additional copy of *ABF2, MIP1,* and *RIM1* | This study | S9 |
| AFY101-2 | *Mat α; ADE2*<br>*ura3::ABF2-URA3*<br>*trp1::MIP1-TRP1*<br>*leu2::RIM1-LEU2* | Haploid WT with an additional copy of *ABF2, MIP1,* and *RIM1* | This study | 1 c, d, e, g;<br>2 a-e, g, i, j;<br>4 a-d, f, g;<br>5 a-c, e-h;<br>S1 c, d, h; |

| | | | | |
|---|---|---|---|---|
| | | | | S2 a, b; S6 c, d; S7; S 10; S12<br>S 13 |
| AFY102-1 | *Mat α; ADE2 ura3::ABF2-URA3 trp1::MIP1-TRP1 leu2::RIM1-LEU2* | Haploid WT with an additional copy of *ABF2, MIP1,* and *RIM1* | This study | 1 c, d, e, g;<br>2 a-e, g, i, j;<br>4 a-d, f, g;<br>5 a-c, e, f;<br>S1 c, d, h;<br>S2 a, b; S6 c, d; S7; S 10; S12<br>S 13 a-c |
| AFY109-1 | *Mat α; ADE2 bmh2::KanMX6* | Haploid *bmh2* deletion strain | This study | 5 g, h;<br>S13 d-f |
| AFY111-1 | *Mat α; ADE2 ura3::ABF2-URA3 trp1::MIP1-TRP1 leu2::RIM1-LEU2 bmh2::KanMX6* | Haploid WT with an additional copy of *ABF2, MIP1* and *RIM1,* and an additional *bmh2* deletion | This study | 5 g, h;<br>S13 d-f |
| AFY115-2 | *Mat α; ADE2 rtg2::KanMX6* | Haploid *rtg2* deletion strain | This study | 5 g, h;<br>S13 d-f |
| AFY117-3 | *Mat α; ADE2 ura3::ABF2-URA3 trp1::MIP1-TRP1 leu2::RIM1-LEU2 rtg2::KanMX6* | Haploid WT with an additional copy of *ABF2, MIP1* and *RIM1,* and additional *rtg2* deletion | This study | 5 g, h;<br>S13 d-f |
| ASY014-1 | *Mat a; ADE2 TRP1 mt-LacO CUP1pr-SU9-2xmNEON-LACI-PGK1pr-SU9-mKATE2:KanMX4* | Haploid LacI-LacO-microscopy strain | Seel et al.[1] | S9 |
| ASY020-1 | *Mat a/α; ADE2/ADE2 URA3/ura3 LEU2/leu2* | Diploid WT | Seel et al.[1] | 1 a, b, d, e, g;<br>2 a-c, e, g, i, j;<br>4 a-d, f, g;<br>5 a-c, e, f;<br>S1 a, b, d, h;<br>S2 a, b; S 3 – 7; S10; S11;<br>S13 a-c |
| ASY033-1 | *Mat a/α; ADE2/ADE2 LEU2/leu2 URA3/ura3 pif1::CglaTRP1/PIF1* | Hemizygous deletion of *pif1* | Seel et al.[1] | 1 b; S 1a |
| ASY046-2 | *Mat a/α; ADE2/ADE2 URA3/ura3 LEU2/leu2* | Double hemizygous | Seel et al.[1] | 1 a, b, e, g;<br>2 e;<br>4 a-d, f, g; |

| | | | | |
|---|---|---|---|---|
| | *mip1::CglaTRP1/MIP1 abf2::clonNAT/ABF2* | deletion of *mip1, and abf2* | | 5 a-c, e, f; S2 a, b; S3-7; S 10; S11; S13 a-b |
| ASY051-2 | *Mat α; ADE2 ura3::ABF2-URA3* | Haploid WT with an additional copy of *ABF2* | Seel et al.[1] | 1 c; S1 c |
| ASY057-3 | *Mat α; ADE2 trp1::MIP1-TRP1* | Haploid WT with an additional copy of *MIP1* | Seel et al.[1] | 1 c; S1 c |
| ASY059-4 | *Mat α; ADE2 ura3::ABF2-URA3 trp1::MIP1-TRP1* | Haploid WT with additional copies of *ABF2,* and *MIP1* | Seel et al.[1] | 1 a, c, e; 2 e; 4 a-d, f, g; 5 a-c, e, f; S1 c; S2 a-c; S3-7; S10; S11; S13 a-b |
| ASY075-1 | *Mat α; ADE2 his3::LexA-ER-AD-TF-HIS3 ura3::LexApr-MIP1-CYC1term-URA3 mip1::clonNAT* | Mip1-inducible strain with additional deletion of endogenous *mip1* | Seel et al.[1] | S1 e |
| FPY015-2 | *Mat α; ADE2 ho::ACT1pr-SU9-KAEDE-HI-NESS-CYC1term-PGK1pr-SU9-mSCARLETI3-CYC1term* | Haploid microscopy HI-NESS strain | Deng et al.[2] | 3 b-d; S8 |
| KSY246-1 | *Mat a/α; ADE2/ADE2 LEU2/leu2 URA3/ura3 hmi1::CglaTRP1/HMI1* | Hemizygous deletion of *hmi1* | Seel et al.[1] | 1 b; S 1a |
| KSY253-1 | *Mat a/α; ADE2/ADE2 LEU2/leu2 URA3/ura3 rpo41::CglaTRP1/RPO41* | Hemizygous deletion of *rpo41* | Seel et al.[1] | 1 b; S 1a |
| KSY254-1 | *Mat a/α; ADE2/ADE2 LEU2/leu2 URA3/ura3 mtf1::CglaTRP1/MTF1* | Hemizygous deletion of *mtf1* | Seel et al.[1] | 1 b; S 1a |
| KSY255-1 | *Mat a/α; ADE2/ADE2 LEU2/leu2 URA3/ura3 mhr1::CglaTRP1/MHR1* | Hemizygous deletion of *mhr1* | Seel et al.[1] | 1 b; S 1a |
| KSY256-1 | *Mat a/α; ADE2/ADE2 LEU2/leu2 URA3/ura3 mgm101::CglaTRP1/MGM101* | Hemizygous deletion of *mgm101* | Seel et al.[1] | 1 b; S 1a |
| KSY257-1 | *Mat a/α; ADE2/ADE2 LEU2/leu2 URA3/ura3 rim1::CglaTRP1/RIM1* | Hemizygous deletion of *rim1* | Seel et al.[1] | 1 b; S 1a |
| MMY116-2c | *Mat α; ADE2* | Haploid WT | Skotheim lab stock | 1 a, c, d, e, g; 2 a-c, e, g, i, j; 4 a-d, f, g; |

| | | | | |
|---|---|---|---|---|
| | | | | 5 a-c, e-h; S1 b-d, h; S2 a, b; S3-7; S10; S12; S13 |
| SAMY013-2 | *Mat α; ADE2 ura3::ABF2-URA3 trp1::MIP1-TRP1 leu2::RIM1-LEU2 ho::ACT1pr-SU9-KAEDE-HI-NESS-CYC1term-PGK1pr-SU9-mSCARLETI3-CYC1term-KanMX* | Haploid microscopy HI-NESS strain with an additional copy of *ABF2, MIP1,* and *RIM1* | This study | 3 b-d; S8 |
| SAMY014-1 | *Mat α; ADE2 ura3::ABF2-URA3 trp1::MIP1-TRP1 leu2::RIM1-LEU2 ho::ACT1pr-SU9-KAEDE-HI-NESS-CYC1term-PGK1pr-Su9-mSCARLETI3-CYC1term-KanMX* | Haploid microscopy HI-NESS strain with an additional copy of *ABF2, MIP1,* and *RIM1* | This study | 3 c-d; S8 |
| SAMY015-1 | *Mat α; ADE2 ura3::ABF2-URA3 trp1::MIP1-TRP1 leu2::RIM1-LEU2 ho::ACT1pr-SU9-KAEDE-HI-NESS-CYC1term-PGK1pr-SU9-mSCARLETI3-CYC1term-KanMX* | Haploid microscopy HI-NESS strain with an additional copy of *ABF2, MIP1,* and *RIM1* | This study | 3 c-d; S8 |
| SAMY016-2 | *Mat a/α; ADE2/ADE2 leu2-3/LEU2 URA3/ura3-1 HO/ho::ACT1pr-SU9-KAEDE-HI-NESS-CYC1term-PGK1pr-SU9-mSCARLETI3-CYC1term-KanMX* | Diploid microscopy HI-NESS strain | This study | 3 b-d; S8 |
| SAMY017-4 | *Mat a/α; ADE2/ADE2 leu2-3/LEU2 URA3/ura3-1 mip1::TRP1/MIP1 abf2::clonNAT/ABF2 rim1::HIS3/RIM1 HO/ho::ACT1pr-SU9-KAEDE-HI-NESS-CYC1term-PGK1pr-SU9-mSCARLETI3-CYC1term-KanMX* | Diploid microscopy HI-NESS strain with hemizygous deletion *of abf2, mip1,* and *rim1* | This study | 3 b-d; S8 |

**Supplementary Table 2: Plasmids used for constructions in this study.**

| Plasmid | Description | Origin |
|---|---|---|
| ASE002-2 | *TRP1-MIP1pr-MIP1-MIP1term* | Seel et al.[1] |
| ASE003-1 | *URA3-ABF2pr-ABF2-ABF2term* | Seel et al.[1] |
| FPE002 | *HO-ACT1pr-SU9-KAEDE-HI-NESS-CYC1term-PGK1pr-SU9-mSCARELTI3-CYC1term* | Deng et al.[2] |
| pAF001G | *LEU2-RIM1pr-RIM1-RIM1term* | This study |
| pAF002G | *HIS3-HMI1pr-HMI1-HMI1term* | This study |
| pAF015-11 | *HIS3-MIP1pr-MIP1-MIP1term* | This study |
| pAF020-1 | *HIS3-MGM101pr-MGM101-MGM101term* | This study |

**Supplementary Table 3: qPCR Primers used in this study.**

| Gene | fwd/reverse | Sequence | Origin |
|---|---|---|---|
| COX2 | Fwd | GTTGATGCTACTCCTGGTAGATT | Seel et al.[1] |
| COX2 | Rev | TTGCATGACCTGTCCCACAC | Seel et al.[1] |
| COX3 | Fwd | TTGAAGCTGTACAACCTACC | Seel et al.[1] |
| COX3 | Rev | CCTGCGATTAAGGCATGATG | Seel et al.[1] |
| ACT1 | Fwd | AGTTGCCCCAGAAGAACACC | Claude et al.[3] |
| ACT1 | Rev | GGACAAAACGGCTTGGATGG | Claude et al.[3] |
| MRX6 | Fwd | CATCCGACGTGGTGCTCTTA | Seel et al.[1] |
| MRX6 | Rev | TCTCATCTCTCCCTCCACCC | Seel et al.[1] |
| RDN18 | Fwd | AACTCACCAGGTCCAGACACAATAAGG | Claude et al.[3] |
| RDN18 | Rev | AAGGTCTCGTTCGTTATCGCAATTAAGC | Claude et al.[3] |
| ABF2 | Fwd | CCAACCTTACGTCCTGCTG | This study |
| ABF2 | Rev | CGTCAAACTCCTTCTTCGCC | This study |
| MIP1 | Fwd | CCATCACAAGCAAGAACGGC | Seel et al.[1] |
| MIP1 | Rev | GTCCCTTTCCAGCTCAACCA | Seel et al.[1] |
| RIM1 | Fwd | GTATATGTTGAAGCAGATG | This study |
| RIM1 | Rev | GCATTTTCTTGGCCCTCAGC | This study |
| HMI1 | Fwd | GGTTCTCTTTGACGGCGGTA | This study |
| HMI1 | Rev | ACACAGTTCATGGGTTGGCT | This study |
| AmpR | Fwd | TTACCAATGCTTAATCAG | This study |
| AmpR | Rev | CCCTTCCGGCTGGCTGGTTTA | This study |
